# Spatially resolved gene regulatory and disease vulnerability map of the adult Macaque cortex

**DOI:** 10.1101/2020.05.14.087601

**Authors:** Ying Lei, Mengnan Cheng, Zihao Li, Zhenkun Zhuang, Liang Wu, Yunong Sun, Lei Han, Zhihao Huang, Yuzhou Wang, Zifei Wang, Liqin Xu, Yue Yuan, Shang Liu, Taotao Pan, Jiarui Xie, Chuanyu Liu, Giacomo Volpe, Carl Ward, Yiwei Lai, Jiangshan Xu, Mingyue Wang, Hao Yu, Haixi Sun, Qichao Yu, Liang Wu, Chunqing Wang, Chi Wai Wong, Wei Liu, Liangzhi Xu, Jingkuan Wei, Dandan Chen, Zhouchun Shang, Guibo Li, Kun Ma, Le Cheng, Fei Ling, Tao Tan, Kai Chen, Bosiljka Tasic, Michael Dean, Weizhi Ji, Huanming Yang, Ying Gu, Miguel A. Esteban, Yuxiang Li, Ao Chen, Yuyu Niu, Hongkui Zeng, Yong Hou, Longqi Liu, Shiping Liu, Xun Xu

**Affiliations:** BGI-Shenzhen, Shenzhen 518083, China; Shenzhen Bay Laboratory, Shenzhen 518000, China; BGI Education Center, University of Chinese Academy of Sciences, Shenzhen 518083, China; School of Biology and Biological Engineering, South China University of Technology, Guangzhou 510006, China; Laboratory of Integrative biology, Guangzhou Institutes of Biomedicine and Health, CAS, Guangzhou 510530, China; Huazhen Biosciences, Guangzhou 510900, China; Yunnan Key Laboratory of Primate Biomedical Research, Institute of Primate Translational Medicine, Kunming University of Science and Technology, Kunming 650500, China; BGI-GenoImmune, BGI-Shenzhen, Wuhan 430074, China; MGI, BGI-Shenzhen, Shenzhen 518083, China; BGI-Yunnan, BGI-Shenzhen, Kunming 650106, China; Allen Institute for Brain Science, Seattle, WA 98109, USA; Cancer and Inflammation Program, National Cancer Institute at Frederick, Building 560, Frederick, MD 21702, USA; Shenzhen Key Laboratory of Single-Cell Omics, BGI-Shenzhen, Shenzhen 518120, China; James D. Watson Institute of Genome Sciences, Hangzhou 310058, China.; Guangdong Provincial Key Laboratory of Genome Read and Write, BGI-Shenzhen, Shenzhen 518120, China; Institute for Stem Cells and Regeneration, Chinese Academy of Sciences, Beijing 100101, China; Faculty of Life Science and Technology, Kunming University of Science and Technology, Kunming, Yunnan 650500, China

## Abstract

Single cell approaches have increased our knowledge about the cell type composition of the non-human primate (NHP), but a detailed characterization of area-specific regulatory features remains outstanding. We generated single-cell chromatin accessibility (single-cell ATAC) and transcriptomic data of 358,237 cells from prefrontal cortex (PFC), primary motor cortex (M1) and primary visual cortex (V1) of adult cynomolgus monkey brain, and integrated this dataset with Stereo-seq (Spatio-Temporal Enhanced REsolution Omics-sequencing) of the corresponding cortical areas to assign topographic information to molecular states. We identified area-specific chromatin accessible sites and their targeted genes, including the cell type-specific transcriptional regulatory network associated with excitatory neurons heterogeneity. We reveal calcium ion transport and axon guidance genes related to specialized functions of PFC and M1, identified the similarities and differences between adult macaque and human oligodendrocyte trajectories, and mapped the genetic variants and gene perturbations of human diseases to NHP cortical cells. This resource establishes a transcriptomic and chromatin accessibility combinatory regulatory landscape at a single-cell and spatially resolved resolution in NHP cortex.

## MAIN TEXT

Cortical organization across primate brains is highly similar, as exemplified by the specialized primary visual cortex and the dorsal and ventral visual streams. With a similar neocortex organization in monkey and human brain, the monkey offers a unique model for studying features of human neurodevelopment and neuropsychiatric diseases^1–3^. The cynomolgus monkey (*Macaca fascicularis*) is one of the most studied non-human primates (NHP) in neuroscience and medicine^4, 5^. Recent advances in transgenesis and genome-editing technologies have led to successful development of new cynomolgus monkey genetic models to study human neurological disorders^6–8^, making this species an excellent experimental NHP model for studying higher order brain function.

Single-cell genomic sequencing enables studies of the underlying diversity and regulatory mechanisms of cortical cells at an unprecedented resolution. Single cell analyses of macaque brain by extension therefore stand to provide an in-depth understanding of the human brain and an opportunity to identify markers and molecular signatures of neuropsychiatric diseases. Earlier single-cell RNA-sequencing studies of prenatal and adult macaque brain identified functionally distinct cortical cell types as well as subtypes across multiple cortical brain areas, and gene expression variations across these cell types^2, 9^. These studies do not capture chromatin states that exert fundamental control of regulating gene expression programs and combining epigenetic analysis with gene expression profiling could therefore yield a complementary understanding of the molecular properties of brain cells. For instance, single-cell-based chromatin states assays can identify cell-type-specific transcriptional regulatory elements and predict potential master transcriptional regulators^10^. Similarly, a survey of chromatin states in bulk tissue from several cerebral cortical regions and hippocampus of the macaque brain revealed region-related chromatin accessibility patterns^11^. However, a systematic characterization of single-cell-based chromatin accessibility of the macaque cortex at a region-specific and single-cell resolution, a requirement for advancing the field, remains outstanding.

Recently, development of spatial transcriptomics technology has made it possible to assign gene expression profiles to spatial coordinates in cortical tissues^12, 13^,. In the present study, we sought to understand how the transcriptional and epigenetic regulatory states differ across functionally distinct cortical areas. To achieve this goal, we performed single nuclei ATAC-seq and RNA-seq, combined with spatial-resolved transcriptomic Stereo-seq (Spatio-Temporal Enhanced REsolution Omics-sequencing) on three functional diverse cortical regions of cynomolgus monkey brain: prefrontal cortex (PFC) and primary visual cortex (V1), two distant areas at frontal lobe and occipital lobe that the differences of neurons between these two areas were recognized in human brain development, and primary motor cortex (M1), a key structure involved in locomotion. Through our massive parallel and integrative topographical analysis, we defined cell type-specific and regional-specific regulatory elements, for the first time and in a spatially resolved manner, resolving single cell gene expression programs between neural cell types, in particular excitatory neurons, in different cortical areas of the NHP brain. We also applied our dataset to delineate the dynamic regulatory landscape of myelination, linking macaque cortical cell types to human neurological disease risk. We provide an interface website (https://db.cngb.org/mba) to present data for exploration and sharing. This publicly available database allows a user to filter cells in the atlas data by brain region, cluster, and cell type, and perform interactive searches from our macaque cortical dataset.

### Single-cell chromatin states define brain cortical cellular taxonomy in adult cynomolgus monkey

We applied modified combinatorial barcoding-assisted single-cell ATAC-seq^10^ and drop-seq based DNBelab C4 ATAC-seq to tissue samples from three cortical areas of the cynomolgus monkey brain: PFC, M1 and V1 (Fig. 1a). To exclude low-quality cells, we filtered snATAC-seq data using a cutoff of 3,000 unique nuclear fragments per cell and a TSS enrichment score of 3 (Extended Data Fig. 1). We then processed a total of 220,040 qualified single cells for further analysis: 72,714 cells from PFC, 70,050 cells from M1 and 77,276 cells from V1 (**Supplementary Table 1**). The cells exhibited a 12,823 median fragment depth per nucleus and a median fraction of reads in peak regions at 55%. We performed iterative latent semantic indexing and batch corrections by ArchR 1.01^14^ with the top 15,000 accessible windows, then used shared nearest neighbor (SNN) clustering by Seurat V3 to separate cells into 23 distinct clusters. (Fig. 1b, Extended Data Fig. 1). Through promoter accessibility and gene activity score, calculated by ArchR of brain cell marker genes, we manually annotated the identity of the cell clusters (Fig. 1c). Using this approach, we characterized seven major cortical cell populations, including excitatory neurons (hereafter labelled as EX, accessible at *NEFH* and *SLC17A7*), inhibitory neurons (IN, accessible at *NEFH* and *GAD1*), oligodendrocytes (OLI, accessible at *MOBP*), oligodendrocyte precursor cells (OPC, accessible at *PDGFRA*), microglia (MIC, accessible at *AIF1*), astrocytes (AST, accessible at *SLC7A10*), endothelial cells (ENDO, accessible at *CLDN5*) (Fig. 1c). The four IN subclusters could be further assigned to VIP, LAMP5, SST and PVALB, respectively, due to the distinct open peaks at the promoter of these genes (Fig. 1c). It’s well known that cortical IN roughly fall into two major branches corresponding to their developmental origins in the caudal ganglionic eminence (CGE) and medial ganglionic eminence (MGE), respectively. *ADARB2*^15^, a marker gene of CGE-derived IN, had accessible chromatin at its promoter in both VIP and LAMP5 sub clusters, while *LHX6*, a marker gene of MGE-derived IN, revealed higher accessibility of its promoter site in SST and PVALB sub clusters. These relationships indicate that, during corticogenesis, VIP and LAMP5 IN are derived from CGE while SST and PVALB are derived from MGE. Notably, we found that 49-56% of differentially accessible sites across all clusters are located at a distance between 10 kb to 100 kb from transcription start sites (TSS), indicating a critical role for distal regulatory elements in specialized cortical cell functions (Extended Data Fig. 1e).

**Figure 1.**
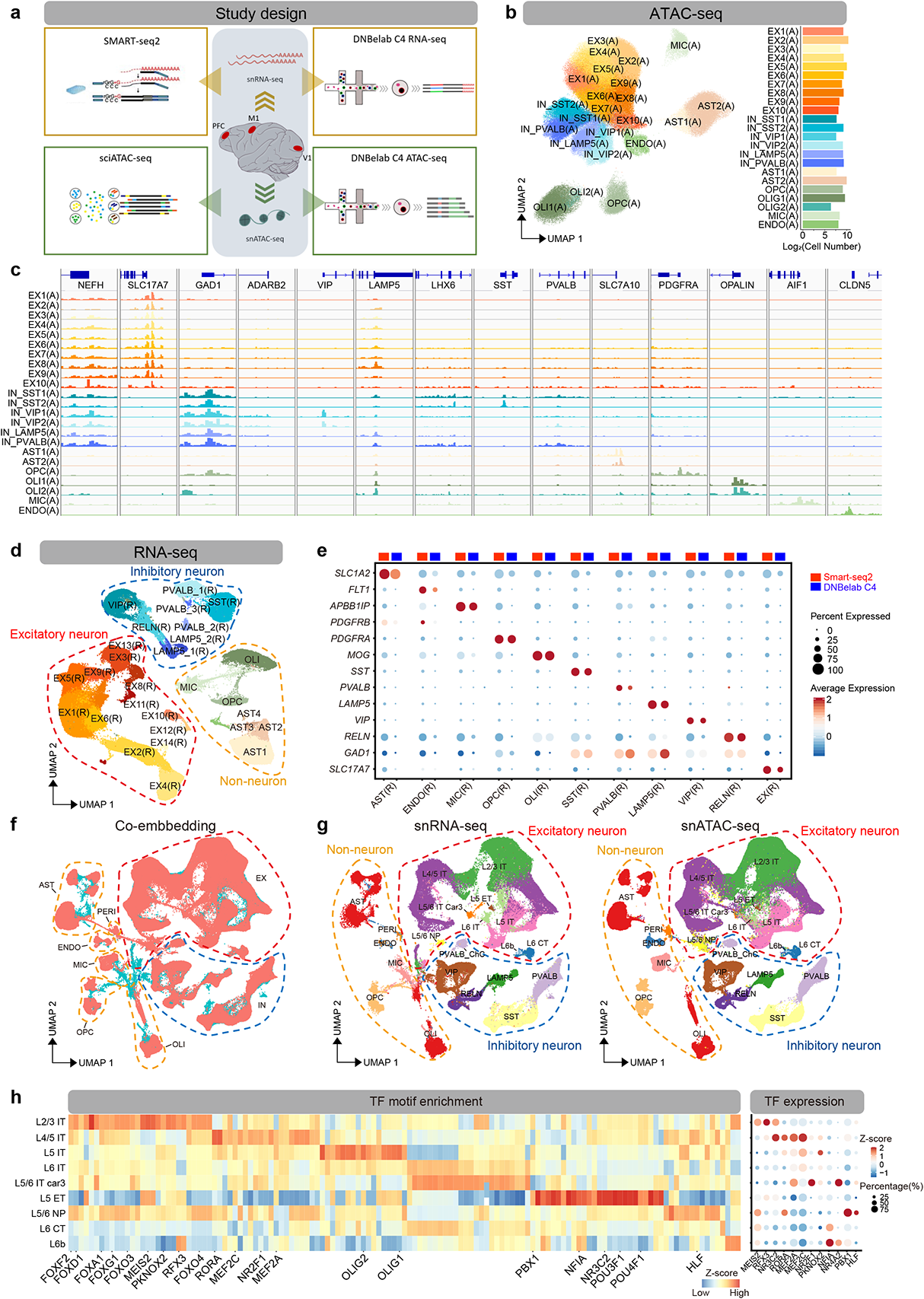
Single nucleus chromatin accessibility and transcriptome of macaque cortex. **a,** Schematic workflow of single-nucleus isolation from the prefrontal cortex (PFC), primary motor cortex (M1) and primary visual cortex (V1) for snATAC-seq and snRNA-seq, sample processing, library generation and downstream analysis. **b,** Uniform manifold approximation and projection (UMAP) visualization of snATAC data showing all clusters colored by cell types (N =220,040 cells) (left), bar plot showing the number of cells per cluster (right). **c,** Integrative genomics viewer (IGV) plots showing color-coded aggregate read density for cells within each cell cluster at neuronal and non-neuronal cell-type-specific marker genes. **d,** UMAP projection of snRNA-seq data showing all clusters colored by cell types (N= 138,197 cells). **e,** Dot plots of expression values for neuronal and non-neuronal cell type-specific marker genes in Smart-seq2 and DNBelab C4 snRNA-seq data. **f,** Co-embedding of snRNA-seq and snATAC-seq clusters across PFC, M1 and V1 brain regions using Seurat V3 (N = 347,682 cells). **g,** UMAP projection of snRNA-seq (N= 138,197 cells) and snATAC-seq cells (N= 209,485 cells) from f). Cell clusters in each UMAP are color-coded and annotated according to cell identity. **h,** Heatmap showing the transcription factor (TF) binding motifs differentially enriched in excitatory neuron subtypes (left), and transcription factors showed differential gene expression in corresponding subtypes of excitatory neuron (right).

To find transcriptional regulators that specify cortical cell type identity, we performed transcription factor (TF) motif enrichment analysis at differentially accessible open chromatin regions in each cell type or subtypes (Extended Data Fig. 1g, **Supplementary Table 2**). This analysis showed that major cell type-enriched TF binding motifs in mouse and human brains^16, 17^ are present in the corresponding macaque cortical cell types. NEUROD1, NEUROG2 and MEF2A, MEF2B and MEF2C motifs were enriched in EX clusters, consistent with their roles in EX specification and activity-dependent regulation on synapse numbers^18, 19^. Likewise, we found that SOX-family motifs (SOX TFs contribute to OLI migration, specification, maintenance of OPC state) were enriched in OLI^20–22^, while PU.1 (a myeloid master regulator that controls microglial development and function) motifs were enriched in MIC^23^. Finally, NF1 and LHX2 (known astrogliogenesis regulators) motifs were enriched in AST^24, 25^, and ETS and forkhead (FOX) transcription factor (endothelial gene expression regulators) motifs were enriched in endothelial cells^26^. These findings demonstrate that regulatory signatures for major cortical cell types are conserved across species. In our analysis, we also identified TF motifs that were enriched in neuron subtypes, such as RORγ, TGIF1/2 and RFX1/2 differentially enriched in EX sub-clusters; MEF2B/MEF2C/MEF2D in parvalbumin (PVALB) INs, ZBTB18 in somatostatin (SST) INs, and ATHB family members in Lysosomal Associated Membrane Protein Family Member 5 (LAMP5) INs.

### Linking chromatin accessibility to transcriptome in monkey cortical cell types

By applying Smart-seq2 based snRNA-seq and drop-seq based DNBelab C4 RNA-seq^27^ to PFC, M1 and V1 of the macaque neocortex, we next assessed how the single-cell epigenetic profiles correlated with the single-cell transcriptomic profiles (Fig. 1a). After quality filtering and cell clustering, we obtained a 11,194 single-nuclei Smart-seq2 transcriptome (6,715 from PFC, 1,943 from M1 and 2,536 from V1) and a 127,003 single-nuclei DNBelab C4 transcriptome (30,844 from PFC, 62,263 from M1 and 33,896 from V1) from which we identified 30 distinct clusters visualized by UMAP (Fig. 1d **and Supplementary Table 3**). Based on known marker genes for cortical cell types, all clusters were annotated as EX, IN and non-neuronal cells, including OLI, OPC, AST, MIC and ENDO (*SLC17A7* for EX, *GAD1* for IN, *MOBP* for OLI, *SLC1A2* for AST, *APBB1IP* for MIC and *FIT1* for ENDO (Fig. 1e, Extended Data Fig. 2; **Supplementary Table 3**). IN subtypes were assigned to CGE-and MGE-derived lineages with mutually exclusive markers. The chandelier cell (ChC) subtype of PVALB-2 inhibitory neuron can further be demarcated by the marker genes UNC5B and RORA^28^ (Extended Data Fig. 2c). By gene ontology (GO) analysis of differentially expressed genes (DEGs), we found that the major cell types were in concordance with the expected corresponding biological processes (Extended Data Fig. 2g), solidifying our cell type assignments.

To understand functional relationships between the transcriptome and open chromatin, we co-embedded the snRNA-seq and snATAC-seq data of cortical cells with Seurat V3^29^. We converted the accessible peak of snATAC-seq data to gene activity score using ArchR and then anchored this analysis to snRNA-seq gene expression data using Seurat V3. With this approach, cells from snATAC-seq were positioned proximal to cells with matching snRNA-seq data assignments (Fig. 1f). Our snATAC-seq data contained similar sub clusters corresponding to the same major brain cell types as the snRNA-seq. Notably, pericytes (PERI) can be distinguished from ENDO cell types both in transcriptomics and in epigenetic nuclei; and the epigenetic LAMP5 IN subtype fell into two sub clusters, one that co-clustered with the transcriptomic LAMP5 subtype, while the other co-clustered with the reelin (RELN) expressing transcriptomic subtype (Fig. 1g). The epigenetic IN PVALB-ChC subtype can be resolved by co-embedding with transcriptomic PVALB-ChC. To further support the concordance of our cell type identification between the snRNA-seq and snATAC-seq datasets, we looked at marker gene expression in the identified transcriptomic cell types and gene activity scores in the corresponding epigenetic cell types (Extended Data Fig. 3).

Given the characteristic neocortical laminar organization and subcortical projections^28, 30^, to further define the subtypes of excitatory neuron, we first adopted the human brain single nuclei RNA-seq dataset from multi-cortical areas (https://portal.brain-map.org/atlases-and-data/rnaseq/human-multiple-cortical-areas-smart-seq) to assign layer and projection pattern onto our transcriptomic excitatory neurons, and then transferred the cell type identification to epigenetic excitatory neurons in the integrated analysis by Seurat V3 (Extended Data Fig. 4a). Thus, the transcriptomic and epigenetic EX subtypes were defined both by the cortical layers (L2/3, L4/5, L5 and L6) and neuronal projection within the cerebral hemispheres, or telencephalon, i.e., intra-telencephalic (IT), extra-telencephalic (ET), near-projecting (NP) and corticothalamic (CT) types (Fig. 1g). To confirm congruence between the transcriptomic and epigenetic EX subtypes, we performed differential gene expression and differential accessible peak tests for layer/projection-defined EX subtypes (Extended Data Fig. 4b and 4c, **Supplementary Table 4**). Of all unique 96,303 differential accessible sites across the 9 subtypes of EX, 4,125 (4.2%) were located within the promoter (TSS ± 1 kb) region. We also examined marker gene expression of mouse cortical EX-subtypes^31^ in our data, whose enriched pattern is highly congruent between macaque and mouse (Extended Data Fig. 4d), providing strong support for stratified EX subtypes. Notably, although the L4 layer is assumed to be absent in M1, an L4-like IT layer was recently identified in human M1, and in mouse, an L4/5 IT layer was also labeled by a combination of L4 and L5 marker expression, such as CUX2, RORB and FEZF2^28, 31^. We confirmed that our L4/5-like IT layer in macaque M1 also expressed CUX2 and RORB and is potentially located between deep L3 and superficial L5 (Extended Data Fig. 4e).

Next, we sought to gain insight into the transcriptional regulatory program in excitatory neurons that underly subtype-specific chromatin accessibility. To this end, we used ChromVAR to analyze TF binding sites enrichment and identify differentially enriched TF binding sites in stratified excitatory neurons. We found that FOX family and MEIS family motifs are significantly enriched in L2/3 IT, that OLIG1/2/3 motifs are strongly enriched in L5 IT whereas POU family motifs are enriched in L5 ET (Fig. 1h). In addition, we found 12 transcription factors that were enriched in EX subtypes as predicted by their binding motifs, including cortical neurogenesis regulation co-factors MEIS2, enriched in L2/3 IT^32^, regional identity, and laminar patterning transcription factor PBX1, in L5/6 NP^33^, enrichment of neurodevelopment regulator NR4A2^34^ and NR2F1^35^ in L5/6 IT Car3, and cortical development transcription factor NFIA^36^ enriched in L6 CT and L6b (Fig. 1h).

In summary, our results demonstrate robust congruence between epigenetic and transcriptomic data for classifying cortical cell types, enabling further integrative analysis.

### Areal heterogeneity of excitatory neuron revealed by integrative single-cell analysis

Recent single-cell studies have resolved area-specific transcriptomic features of cortical EX in mice^30^, and the molecular and epigenetic signatures in the developing human brain^37, 38^. However, the epigenetic features that correspond to area-specific feature in adult cortex (monkey or human) are yet to be clarified. We were therefore intrigued to see area-specific transcriptomic and epigenetic clusters of excitatory neurons in the frontal cortex (referring to PFC and M1) and V1, while no such cortical area-separated clusters were identified in IN and non-neuronal cells (Extended Data Fig. 5a).

We performed an inter-regional comparison of the transcriptomic features across PFC, M1 and V1 using drop-seq based DNBelab C4 RNA-seq cells (127,003 cells), and identified 699, 125 and 388 genes that were area-specifically enriched in EX, IN and non-neuronal cell types/subtypes, respectively (**Supplementary Table 5**, Extended Data Fig. 5b-5d). In EX, some area-enriched genes are broadly upregulated in cortical layers, such as *SNTG2* in L2/3 IT, L4/5 IT, L5 IT and L6 IT of M1, while some genes are enriched in specific layers, such as *KCNH8* in L4/5 IT of V1 (Extended Data Fig. 5e). We found genes differentially expressed in PFC and V1 areas of adult macaque bulk tissue^39^ are expressed in EX-subtypes of corresponding layers, including *DPYD*, which is upregulated in PFC EX subtypes, and *ZFPM2*, *MAML3* and *VAV3*, which are upregulated in V1 EX subtypes (Extended Data Fig. 5e). Moreover, genes reported as differentially expressed in human excitatory neurons of PFC versus V1^40, 41^ were similarly enriched in macaque, including *L3MBTL4* and *TLL1,* which were upregulated in PFC; and *PDE3A*, *TRPC3* and *PCDH7*, which were upregulated in V1 (Extended Data Fig. 5e), indicating a conserved area specialization between human and macaque cortex. Of note, we not only identified layer and neuron projection specificity in previous known area-enriched genes for PFC or V1, but also identified new area-specific candidate EX genes, such as *GALNTL6* and *B4GALNT3* in PFC, *SNTG2* and *PRUNE2* in M1, and *KCNH8*, *HPSE2*, and *ADAMTS17* in V1 (Extended Data Fig. 5e).

To identify area-specific chromatin opening states, we performed an inter-regional comparison of chromatin accessibility using the snATAC-seq cells with matched cell type identification from snATAC-seq annotation and by snRNA-seq transferring (209,485 cells) and found 83,110 differential accessible sites (DA peaks) between PFC, M1 and V1, across 9 subtypes of EX (**Supplementary Table 6**). Among of them, 6,347 (7.64%), 15,885(19.11%) and 42,528 (51.17%) of differential accessible sites were located within 0 to 1 kb, 1 kb to 10 kb and 10 kb to 100 kb genomic distance to the nearest genes, respectively. Using the matched cell type of single cell transcriptomic and chromatin accessibility data, we built the links of the *cis*-regulatory elements (CREs) to targeted gene expression by ArchR in each cell type/subtype (see methods). In total, we identified 54548 significant associated CRE-gene pairs, with a median of 3 CREs linked to each of 12484 genes, similar as reported in human brain cell types^42^ (Fig. 2a), and most of the gene-linked CREs showed a cell type-specific distribution (Fig. 2b). For all neuron and non-neuronal cell type/subtypes, we found a significant overlap between CRE-targeted genes and cell type/subtype DEGs (Extended Data Fig. 6), demonstrating a key role of chromatin accessible sites in defining cell type specific transcriptomics. Next, we sought to elucidate the peak-gene regulation on areal heterogeneity. For most excitatory neuron subtypes, there is a significant overlap between DA peak-targeted genes and the differentially expressed genes among PFC, M1 and V1 (Fig. 2c), indicating an important role of differentially accessible sites in area-specialized transcriptomics.

**Figure 2.**
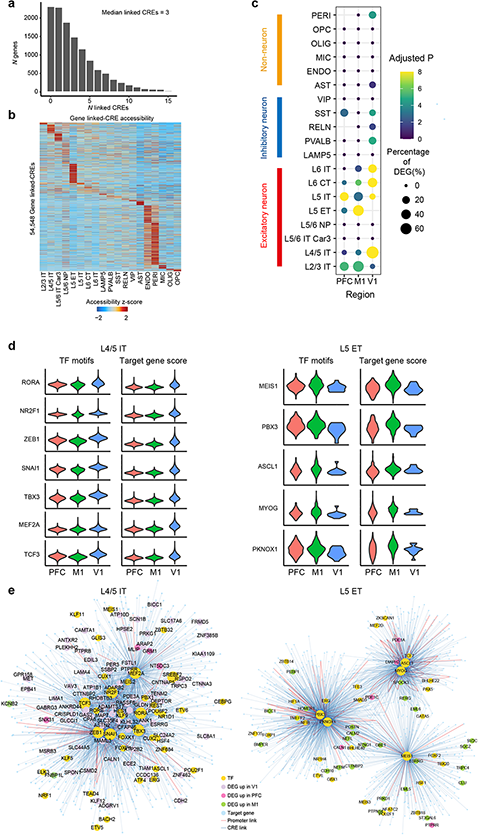
Transcriptional regulation of the cell type-specific and area-specific gene expression. **a,** Histogram showing the number of genes that linked to *cis*-regulatory elements (CREs). **b,** Heatmap showing row-normalized gene-linked CREs in each cell type/subtype of snATAC-seq cells. **c,** Dot plots of overlapped genes between differentially accessible peaks (DA peaks)-targeted gene and the differentially expressed genes (DEGs) of each cortical area in neuronal and non-neuronal cell types/subtypes. Dot size represents the percentage of overlap genes of area DEGs. **d,** Violin plots of motif enrichment and targeted gene score of selected TFs that had significant upregulated in V1 of snATAC-seq L4/5 IT type excitatory neuron (left) and in M1 of snATAC-seq L5 ET type excitatory neuron (right). **e,** TF regulatory networks showing the predicted candidate target genes for transcription factors RORA, NR2F1, ZEB1, SNAI1, TBX3, MEF2A and TCF3 in L4/5 IT type excitatory neuron (left) and the predicted candidate target genes for transcription factors MEIS1, PBX3, ASCL1, MYOG and PKNOX1 in L5 ET type excitatory neuron (right).

To further explore the transcriptional regulation of the area-specific expression, we first measure transcription factor binding site enrichment in each EX-subtypes using ChromVAR and analyzed the differentially enriched TF binding motif across PFC, M1 and V1. We showcase transcription factors in L4/5 IT and L5 ET, we found the binding motif enrichment and significantly increased targeted gene score of ZEB1, MEF2A, RORA, NR2F1, SNAI1, TBX3 and TCF3 in V1 of L4/5 IT, some of these have previously been reported for motif enrichment in human developing V1^37^, and MEIS1, PBX3, ASCL1, MYOG and PKNOX1 in M1 of L5 ET (Fig. 2d). Then, we constructed the TFs regulated genes network using peak-gene links and TF binding motif, the area-specific DEGs were identified among the TFs targeted genes (Fig. 2e). Collectively, our findings reveal overlap in inter-regional transcriptomic profiles and chromatin accessibility and identified transcriptional regulation in areal heterogeneity of excitatory neuron.

### Specialized transcriptomics in macaque prefrontal cortex and motor cortex

Areal and layer proximity is the strongest determinant for gene expression relationships in the primate cortex^1^, which is believed to explain why V1 has a more distinct transcription pattern when compared to PFC and M1. Previous studies have revealed fundamental differences between the anterior and posterior cerebral cortex by comparing differentially expressed genes between PFC and V1, whereas heterogeneity within the frontal cortex of primates is less well understood. Here, we analyzed differentially expressed genes between PFC and M1 in excitatory neuronal subtypes and found that 210 genes and 291 genes were upregulated in PFC and M1 across EX subtypes, respectively. (Fig. 3a, **Supplementary Table 7**). Specialized functions and expressions have been reported in L5 extratelencephalic (L5 ET) neurons of primate and mouse motor cortex^28^. In M1 L5 ET, we identified many genes involved in calcium ion transport, such as *SLC24A2*, *ITPR2*, *CALM2*, *PDE4B* and *CAMK2G* (Fig. 3b **and** Fig. 3f). In our data sets, we didn’t find reported primate M1 L5 ET-enriched (versus M1 L5 IT type) potassium ion signaling-related genes, such as *KCNN2*, *KCNA1* and *HCN1*^28^. To ascertain PFC- and M1-associated specialized biological functions in EX subtypes, we performed gene ontology analysis. We found that many M1-enriched genes are involved in ‘muscle contraction’ and comprise genes encoding for the adrenergic receptor and calcium ion transport proteins, exemplified by L6 CT type, (Fig. 3c and 3f), suggestive of an involvement in the long-range cortico-motor neuronal projections^43^. For PFC-enriched genes in EX subtypes, we found glutamate receptor signaling pathway genes to be significantly increased, including *GRIK1*, *GRIK2*, *GRID2*, *GRIK1*, *GRIK4* and *GRIN2B,* exemplified by L5 ET type, (Fig. 3d and 3f). We also found enrichment of many axon guidance genes implicated in cortical development and psychiatric disorders ^44–46^, such as *SLIT/ROBO* signaling members *SLIT1*, *SLIT3* and *ROBO1* and semaphorins *SEMA3E*, *SEMA4A, SEMA3D* and *SEMA6D*, exemplified by L4/5 IT, L6 IT and L6b type (Fig. 3e and 3f). These axon guidance genes might influence distinct PFC connectivity patterns.

**Figure 3.**
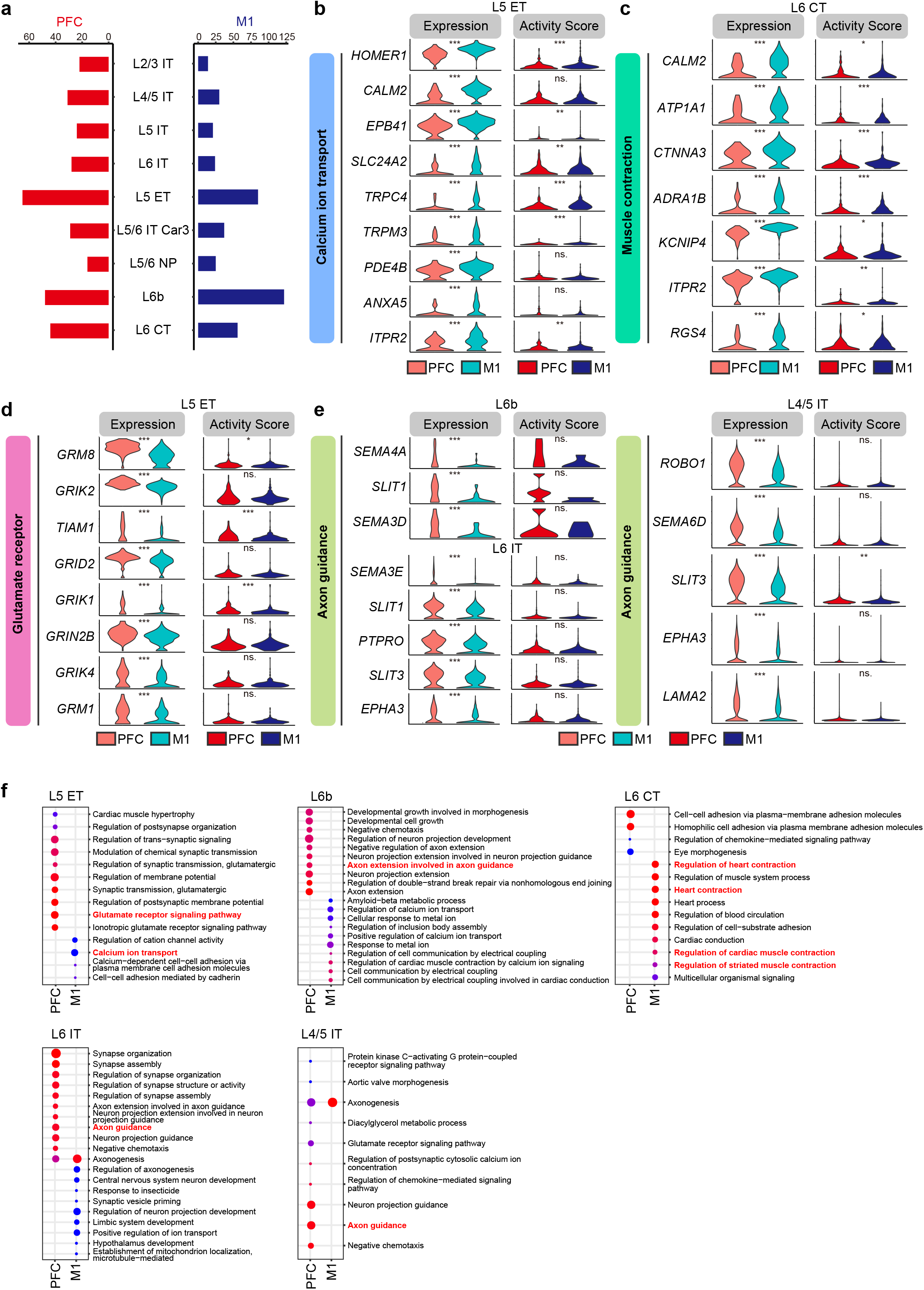
Specialized transcriptomics of excitatory neuron between macaque prefrontal cortex and Motor cortex. **a,** Number of differentially expressed genes between PFC and M1 in each subtype of excitatory neurons. **b,** Violin plots of calcium ion transport-related gene expression (left) and gene score (right) that are significant enriched in L5 ET type excitatory neuron of M1. \**P* < 0.05, \*\**P* < 0.01, \*\*\**P* < 0.001. **c,** Violin plots of ‘muscle contraction’-related gene expression (left) and gene score (right) that are enriched in L6 CT type excitatory neuron of M1. \**P* < 0.05, \*\**P* < 0.01, \*\*\**P* < 0.001. **d,** Violin plots of glutamate receptor signaling-related gene expression (left) and gene score (right) that are enriched in L5 ET type excitatory neuron of PFC. \**P* < 0.05, \*\**P* < 0.01, \*\*\**P* < 0.001. **e,** Violin plots of axon guidance-related gene expression and gene score that are enriched in L4/5 IT, L6 IT and L6b type excitatory neuron of PFC. \**P* < 0.05, \*\**P* < 0.01, \*\*\**P* < 0.001. **f,** Gene ontology terms enriched among differentially expressed genes of PFC and M1 in L5 ET type, L4/5 IT, L6 IT, L6 CT and L6b type excitatory neuron.

### Gradient gene expression pattern of excitatory neuron across laminar and cortical regions

Tissue level transcriptomics in macaque brain^2, 39^ and single cell transcriptomics in multi-regions of mouse cortex ^31^ suggest that a continuous gradient expression likely drives many of the adult cortex areal expression differences. Here, we sought to further reveal two aspects of the inter-areal diversity of excitatory neurons: 1) The gradient gene expression patterns across cortical layers; and 2) The gradient gene expression patterns across PFC, M1 and V1 areas. We first analyzed the gene expression pattern in the largest branch of excitatory neurons; the inter-telencephalon (IT) projection type (corresponded well with cortical depth), and clustered these genes into 5 gradient expression patterns across L2/3 IT, L4/5 IT, L5 IT and L6 IT using the soft clustering package Mfuzz^47^ in PFC, M1 and V1, respectively (Extended Data Fig. 7a and 7b) to reveal gene distribution and enrichment in specific layer of cortex. Then, we compared the gene sets in consensus patterns between PFC, M1 and V1. We found that 119 genes were uniformly expressed across layers, including transcriptional factors *MEIS2*, *RFX3*, *HIVEP1*, *ARID2* and *PKNOX2*, and glutamate receptor *GRIA4* and *GRIN2B*; while 1694 genes showed distinct expression pattern between PFC, M1 and V1, and were enriched in biological processes such as cell morphogenesis, synaptic signaling and regulation of cell projection organization (Extended Data Fig. 7c, **Supplementary Table 8**).

When we analyzed the gradient gene expression patterns across PFC, M1 and V1 areas, we obtained 6 clustered patterns in each excitatory neuron subtype (Extended Data Fig. 7d and 7e, **Supplementary Table 9**). These patterns reflect the cortical area-preferred gene expression, for example, pattern 1 and pattern 5 are gradient expression for PFC-preferred and V1-preferred genes, respectively. Notably, the gradient expression pattern of the axon guidance molecule *SLIT1* (pattern 1) is highly consistent with previous *in situ* hybridization analysis in macaque cortex; highest in the prefrontal cortex, faint in the primary motor cortex and lowest in the primary visual cortex^44^, in congruence with the gradient expression pattern of macaque cortical genes predicted by our dataset. When we examined the consensus gene patterns in different EX subtypes, for example, genes in pattern 1 of L2/3 IT (163 genes), L4/5 IT (192 genes), L5 IT (203 genes) and L6 IT (177 genes), we found that only 6 genes were shared by all four subtypes and that 126 genes were shared by more than one subtype, including the *TCF4*^48^, a TF implicated in schizophrenia, which was shared by L2/3, L4/5 and L6. Moreover, genes in pattern 1 are involved in divergent signaling pathways, for instance GTPase activity regulating genes are enriched in L2/3, genes involved in regulation of macro-autophagy are enriched in L4/5 and genes involved in membrane docking are enriched in L6 (Extended Data Fig. 7f). These findings demonstrated the cell subtype/laminar-biased pattern for different cortical functions.

### Spatial gene expression characterized by Stereo-seq

To spatially resolve the macaque cortical cell transcriptome, we performed the recently developed technology Stereo-seq^49^, in which DNA nanoball (DNB) patterned array chips are combined with *in situ* RNA capture to enable nanoscale transcriptome analysis of tissue sections. With the DNB approach, Stereo-seq achieves a resolution of 500 or 715 nm and captures a significantly higher number of spots per 100 μm^2^ compared to other related techniques^49^. We used 10 μm paired adjacent tissue sections from macaque PFC, M1 and V1 tissues for Nissl staining or Stereo-seq, respectively (Fig. 4a). Stereo-seq captured an average number of 31,916 of 37.5 μm bins (bin 50, 50 × 50 DNB) per section with 1,748 genes and 4,754 transcripts per 37.5 μm bin for analyzed sections (Extended Data Fig. 8). We performed unsupervised clustering with gene matrices of 37.5 μm bin with Seurat V3 and found that the clusters defined by the Stereo-seq data shows high similarity with cortical layers defined by Nissl staining in adjacent tissue sections, as exemplified by recognition of the L4 multi-sub-layers in V1 (Extended Data Fig. 9a). Thus, the high resolution achieved with Stereo-seq data enabling us to distinguish heterogenous features of macaque cortex tissue.

**Figure 4.**
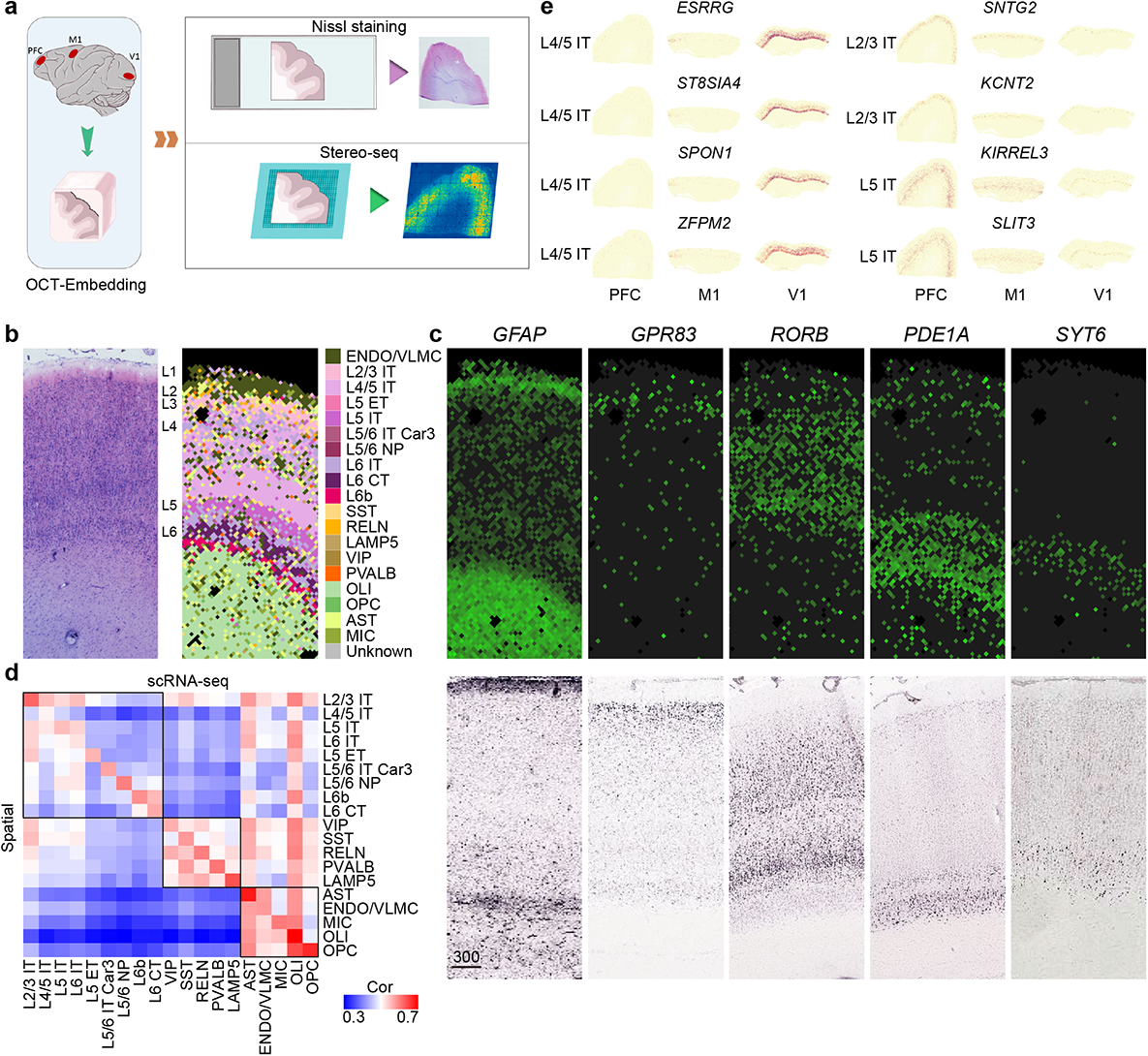
Characterization of cortical gene expression by Stereo-seq. **a,** Schematic workflow of tissue sampling and Stereo-seq from the prefrontal cortex (PFC), primary motor cortex (M1) and primary visual cortex (V1). **b,** Nissl staining of adjacent section to stereo-seq section of primary visual cortex (left), and cell type/subtypes annotation for 37.5 μm bins of stereo-seq section of primary visual cortex by snRNA-seq data (right). **c,** Expression of known layer-marker genes in the stereo-seq section of b) (top), and RNA ISH images for corresponding layer markers in visual cortex of adult macaque brain from NIH Blueprint NHP atlas (https://www.blueprintnhpatlas.org/) (bottom). **d,** Heatmap showing Spearman’s rank correlation between clusters identified in Stereo-seq versus snRNA-seq data. **e,** Spatial expression maps of selected differentially expressed genes between PFC, M1 and V1 in EX subtypes.

To register the spatial localization of macaque cortical cell types, we assigned the cell type/subtype from our snRNA-seq data to Stereo-seq data using SPOTlight^50^. We found that the cell type annotations were highly concordant with spatial expression of known layer markers from macaque brain^1^ and the cortical layers defined from histology staining of adjacent tissue section (Fig. 4b-d, Extended Data Figure 9b). We then examined the area-differentially expressed genes of excitatory neuron subtypes between PFC, M1 and V1 of snRNA-seq in the corresponding cell subtypes annotated in Stereo-seq sections of PFC, M1 and V1. We found a significant number of genes were also exhibit area-specificity in Stereo-seq data (42 genes in PFC, *P* = 5.13 x 10^-9^; 51 genes in M1, *P* = 3.10 x 10^-5^; 81 genes in V1, *P* = 0.025, by hypergeometric test), including known and newly identified area-enriched genes in EX (Fig. 4e, **Supplementary Table 10**). demonstrating the consistent finding for area-specific transcriptomics in cell subtype resolution between single cell transcriptomic and Stereo-seq.

Next, to explore the gradient gene expression across cortical layers *in situ*, we examined the spatial distribution of genes with gradient gene expression patterns of snRNA-seq (see methods). For the 41 genes showed gradual decreased expression pattern from upper to lower layer in PFC, M1 and V1 of snRNA-seq cells (pattern 1 in Extended data Fig. 7a and 7b, Supplementary Table 8), we found 9 genes showed gradual decreased expression across upper to lower layer in Stereo-seq sections of PFC, M1 and V1, including glutamate receptor *GIRA4*, schizophrenia risk gene *ARHGAP10*^51^ and important transcription factor of central nerve system *RFX3*^52^ (Extended Data Fig. 10).

Collectively, we demonstrated that integrative analysis of single cell genomics and spatial resolved transcriptomics can identify consistent areal diversity. Additionally, for the first time, we captured consensus gradient expression pattern in different cortical regions *in situ*.

### Dynamic single cell regulatory landscape of oligodendrocyte trajectory

OLI wrap neuronal axons with myelin to support neuronal function^53^. Previous work has reported on transcriptional and epigenetic regulation pathways of OLI maturation and myelination in mouse models and the human brain^54^, but due to insufficient data coverage it wasn’t possible to assess how these two layers of regulation are dynamically correlated. Given that macaque models are widely used to model disease, such as demyelination-related diseases autoimmune encephalomyelitis and multiple sclerosis^55–57^, we first sought to investigate whether our data could recapture the dynamic signature of OPC differentiation and OLI development in those studies and extend these to human biology, and second whether we could deepen our understanding of the OLI regulatory landscape (Fig. 5a).

**Figure 5.**
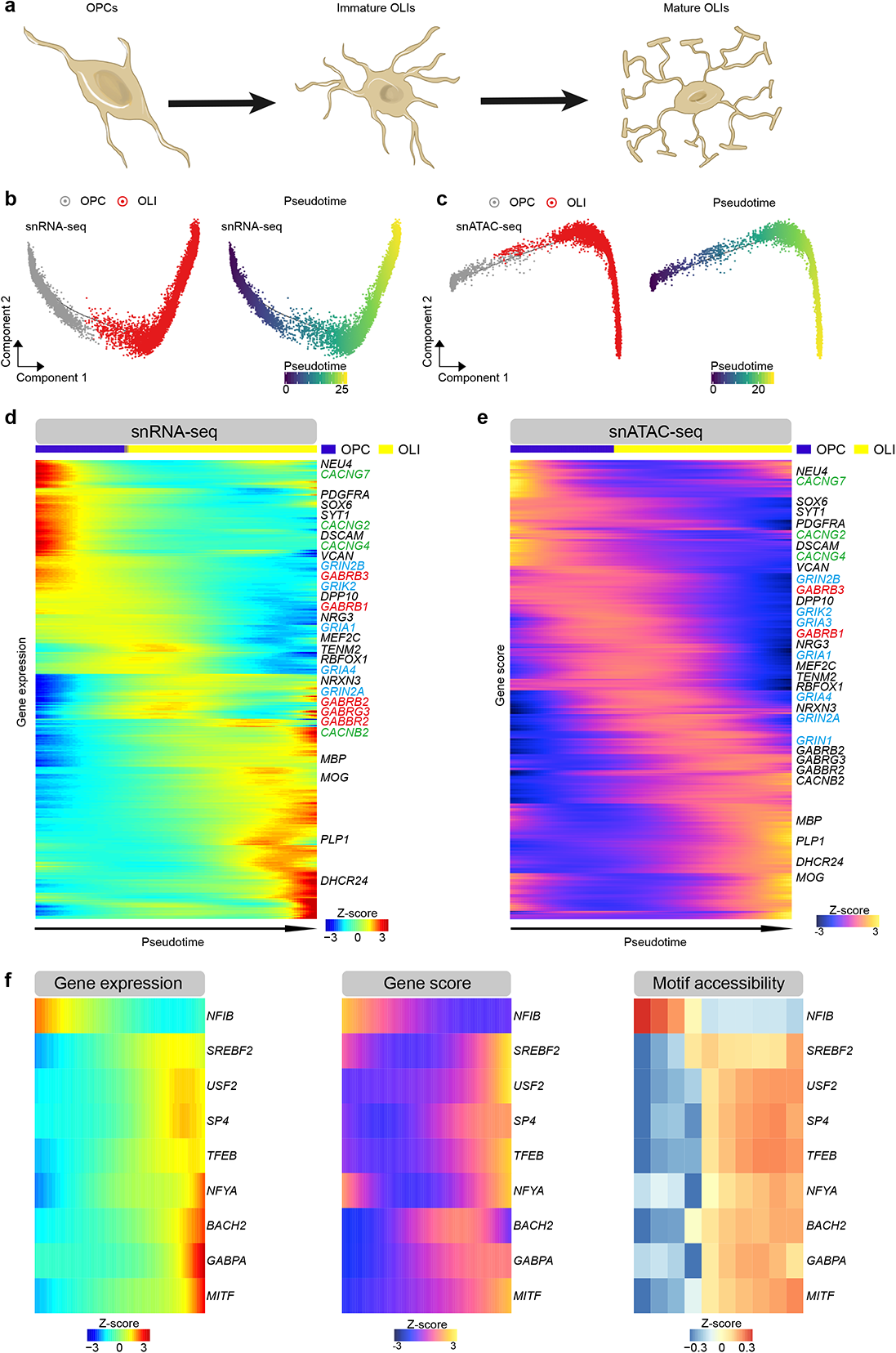
Integrated single cell regulatory landscape of oligodendrocyte trajectory. **a,** Schematic illustration for oligodendrocyte maturation. **b,** Pseudotime trajectory of OPC and OLI indicating the gene expression obtained from snRNA-seq data. **c,** Pseudotime trajectory of OPC and OLI indicating the gene activity score obtained from snATAC-seq data. **d,** Heatmap showing genes expression dynamics across oligodendrocyte lineage pseudotime trajectory indicated in b). Genes of glutamate receptors, gamma-aminobutyric acid (GABA) receptors and voltage-gated calcium channel (VGCCs) were blue, red, and green color-coded, respectively. **e,** Heatmap showing accessibility dynamics (gene activity score) across oligodendrocyte lineage pseudotime trajectory in c). Genes of glutamate receptors, gamma-aminobutyric acid (GABA) receptors and voltage-gated calcium channel (VGCCs) were blue, red, and green color coded, respectively. **f,** Gene expression (left), gene activity score (middle) and motif enrichment (right) of 9 TFs across OLI pseudotime trajectory.

We performed pseudotime ordering of cells by Monocle 2 to construct OLI trajectories using gene expression and gene activity score (Fig. 5b and 5c). We found that both the marker gene expression and activity score reported in mouse and human OPC and mature OLI^58–60^ were also enriched in our macaque OPC (*NEU4* and *PDGFRA*) and OLI (*MBP* and *PLP1*) (Fig. 5d and 5e). Despite recent studies highlighting the importance of glutamatergic, GABAergic and calcium signaling in the OLI lineage, these relationships remain controversial given limited insight into tissue heterogeneity and receptor activation during OLI development and maturation. In earlier work, GO enrichment of glutamate receptors were confirmed in human OPC during lineage maturation^60^. Here, we found enriched gene expression and high epigenetic score of ionotropic glutamate receptors in macaque OPC and immature OLI, including the NMDA (N-methyl-D-aspartate) receptors (*GRIN2A* and *GRIN2B*) and kainate receptors (*GRIK1, GRIK2, GRIK4* and *GRIK5*) in OPC and immature OLI, AMPA (α-amino-3-hydroxy-5-methyl-4-isoxazolepropionic acid) receptors (*GRIA1*, *GRIA3* and *GRIA4*) in immature OLI. Furthermore, we found enriched gene expression and high epigenetic score of GABA (gamma-aminobutyric acid) receptors (*GABRB1, GABRB2, GABRB3, GABBR2* and *GABRG3*) and voltage-gated calcium channels (*CACNB2, CACNB4, CACNG2*, *CACNG4, CACNG7* and *CACNG8*) in OPC and immature OLI. When connecting the expression trajectories to accessibility dynamics, we defined a concordant set of 2211 genes in expression-activity score pairs (Spearman’s correlation coefficient r_s_ >0.2, *P* < 0.01, Extended Data Fig. 11a,), including transcription factor SOX5, SOX6, OLIG2 and MYRF, known for roles in OPC differentiation and OLI development^54^. GO terms for concordance gene sets supported stage-enriched pathways and dynamic changes of neuronal activity along the pseudotime trajectory (Extended Data Fig. 11a). Previous work proposed that the Wnt signaling pathway plays an important role as a stage-specific and multi-functional regulator of OLI development^61^. Consistent with this study, our data showed that genes involved in Wnt signaling are enriched in OPCs (Extended Data Fig. 11b).

To understand transcriptional regulation of OLI maturation, we characterized TF motif enrichment across OPC differentiation and OLI development by mapping chromVAR TF deviation scores to cells along the pseudo time trajectory. This approach identified 129 TFs that defined different stages of the oligodendrocyte maturation process (Extended Data Fig. 12). To link TF expression to enrichment of TF binding motifs across different stages of OLI maturation, we found 9 TFs that had congruent gene expression and gene activity scores/motif accessibility profiles (Fig. 5f), such as OLI differentiating activators and myelination regulators TFEB in mature OLI^62^.

To gain insight into the similarity and differences of dynamic trajectory of oligodendrocyte in adult macaque monkey and human, we adopted the snRNA-seq and snATAC-seq data of OPC and OLI of adult human brain from a recent study^42^. After constructing the pseudotime trajectory of gene expression and gene activity score, we found a concordant set of 1756 genes in expression-activity score pairs across the human OLI trajectories (Spearman’s correlation coefficient r_s_ > 0.2, *P* < 0.01, Extended data Fig. 13a). Similar with macaque OLI, these sequential activated genes were strongly enriched in trans−synaptic signaling and cellular morphogenesis pathways (Extended data Fig. 13a). Among of them, 829 genes were overlapped with the concordant gene expression-activity score pairs along macaque monkey OLI trajectory (**Supplementary Table 11**), including myelin forming OLI signature MOG, PLP1 and OPALIN, and mature OLI signature KLK6 and SLC5A11^42^. We found 5 transcription factors had congruent gene expression and gene activity scores /motif accessibility profiles along the human OLI trajectory (Extended data Fig. 13b). Of note, activation of transcription factor EB (TFEB) were observed along both human and macaque monkey OLI trajectory; and the sterol regulatory element-binding transcription factor (SREBP), involved in cholesterol and fatty acids biosynthesis of glia cell and associated with schizophrenia^63^, were found activation in mature stage of macaque and human OLI trajectory.

Taken together, our analysis proposes master regulators for OLI maturation based on combinatorial analysis of single nucleus transcriptomic and chromatin accessibility, providing candidates for further functional studies in demyelination diseases and demonstrate the conserved regulatory landscape of oligodendrocyte trajectory in adult human and macaque monkey brains.

### Mapping human disease risk loci to cortical cell types

Linking cell-type specific regulatory elements with disease risk variants identified in genome-wide association studies (GWAS) can help us understand whether specific cell types contribute to disease pathobiology, which in turn can become instrumental for developing targeted therapeutic approaches. Single-cell epigenetic data of mouse and human brain have already proven to be useful for mapping neuropsychiatric disease risks with specific cellular subtypes^60, 64^. To demonstrate the robustness of our dataset, we assessed enrichment of human neurological and neuropsychiatric disease risk factors in macaque cortical cell types. We mapped differentially accessible peaks of each cluster and all accessible peaks to orthologous coordinates in the human hg19 genome, then performed linkage disequilibrium score regression (LDSC) to measure SNP heritability enrichment for human traits within differential accessible peaks of each epigenetic cluster in PFC, M1 and V1, respectively. We adopted the GWAS summary for neurological and psychiatric disorders and neurobehavioral traits from recent studies (**Supplementary Table 12**) as well as non-brain-related diseases (from UK biobank).

Consistent with cell type mapping studies of neurological trait risk in human and mouse, we found a highly significant enrichment of heritability in neurons for neuropsychiatric traits such as major depressive disorders (MDD) and schizophrenia (SCZ) (Fig. 6a)^60, 64^. In line with studies reporting on microglial activation in AD and early AD-associated transcriptional changes occurring in other brain cell types, we found strong enrichment of Alzheimer’s disease (AD) SNP-heritability in MIC^65, 66^. Extending from earlier single cell transcriptomic analysis reporting on enrichment of SCZ genetic variants in cortical pyramidal neuron and interneurons^67^, we here mapped significant enrichment of SCZ genetic variant to excitatory neuron of the L2/3 IT type and SST inhibitory neuron in PFC and M1. The MDD heritability enrichment in prefrontal cortex has been highlighted ^68^, and we discovered here that genetic variants of MDD were enriched in both PFC and M1, but not in V1. Conversely, the SNP-heritability of autism disorder (ASD) was significantly enriched in excitatory neuron of M1 and V1, but not PFC. Consistent with cell types known to be affected in non-brain-related autoimmune diseases, we found SNP-heritability enrichment for asthma, hypothyroidism, and eczema in microglia. Thus, our macaque cortex single-cell open chromatin landscape provides cell type-specific datasets as a resource for evaluating genomic loci implicated in human neurological traits in specific macaque brain cell types and cortical areas.

**Figure 6.**
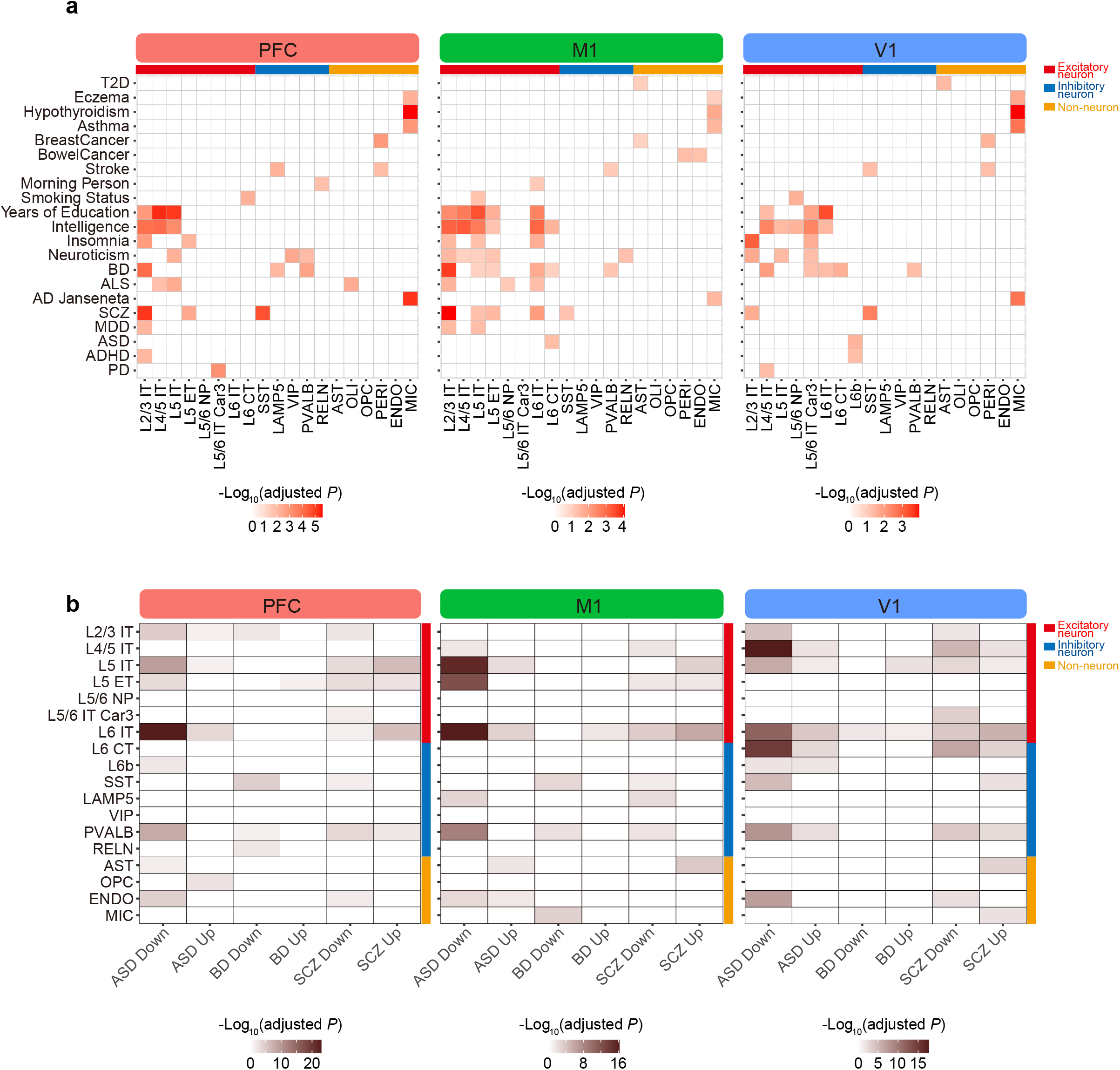
Mapping of disease genes to specific brain cell types. **a,** Heatmap showing the enrichments of genetic variants of human traits among the neuronal and non-neuronal cell types/subtypes from snATAC-seq data of PFC, M1 and V1. AD, Alzheimer’s disease; ADHD, attention deficit hyperactivity disorder; ASD, autism spectrum disorder; BD, bipolar disorder; ALS, amyotrophic lateral sclerosis; MDD, major depression disorders; SCZ, schizophrenia, T2D, type 2 diabetes. **b,** Cell type-enrichment of neurological diseases-affected genes in Stereo-seq sections of PFC, M1 and V1. DE: differential expressed genes (up: up-regulated, down: down-regulated) from brains of individuals with ASD, BD or SCZ compared with neurotypical controls, respectively.

To map spatial enrichment of disease-affected genes in specific cell types, we integrated large scale postmortem brain datasets surveying differentially expressed gene in patients of ASD, bipolar disorder (BD) and SCZ^69^ with our Stereo-seq macaque cortical tissue dataset (Fig. 6b). In neurons, we found extensive enrichment of ASD and SCZ DEGs, but only a mild enrichment for BD DEGs in neuron and non-neuronal cells. We found enrichment of genes downregulated in ASD in L5 IT and L6 IT of PFC, which were also identified as recurrently affected cell types across ASD patients^70^, and observed distinct enrichment of ASD DEGs across PFC, M1 and V1. Further, we found enrichment of SCZ DEGs in deep layer IT neurons and the differential enrichment of up-regulated and down-regulated genes of SCZ in upper layer IT neurons., highlighting SCZ-associated changes on inter-telencephalic projections neurons across layers. Consistent with a recent single cell transcriptomics analysis of SCZ patient-derived brain samples^71^, we found a broad enrichment of SCZ DEGs in excitatory neuron subtypes, with the PVALB-type being the most affected inhibitory neurons. These findings highlight the value of our data for mapping and predicting cell types affected in disease.

A recent study comparing single cell transcriptomes of post-mortem prefrontal cortex obtained from control and SCZ patients suggest a strong correlation between genetic risk loci and transcription perturbations within neuronal cell types^71^. Our results revealed a similar trend for SCZ-associated genes in the PFC region, and specifically that both SCZ-heritability and SCZ-affected genes are enriched in L2/3 IT and SST neuronal cell types. However, this is not the case for ASD; although ASD-affected genes are broadly enriched in neuronal subpopulation in PFC, ASD-heritability is not enriched in any particular cell type in PFC. By combining epigenetic and spatial transcriptomics analysis, we have generated a resource that may provide clinical insights in cell type and cortical region-specific programs underlying genetic and disease states.

## DISCUSSION

Developing therapeutic approaches for central nervous system disorders has been hampered by the lack of models that can adequately mimic human disease pathogenesis. While murine models have been instrumental to study developmental trajectories and the genetic basis of disease phenotypes, they poorly mirror human physiology, behavior, and progressive disease mechanism. Using larger animal models that are phylogenetically close to humans, such as pigs and monkeys is therefore warranted for certain conditions. Despite evolutionary differences in cognitive functions and behaviors, monkeys have the social complexity, brain structure and neuronal circuitries that are more closely related to humans^72^. As such, monkey models faithfully recapitulate several features of many neurodegenerative diseases such as Parkinson’s^73^ and Huntington’s^74, 75^. Therefore, dissecting brain cellular composition and regulatory circuitries at a single cell resolution as well as determining disease-specific molecular signatures in species phylogenetically closer to human is an important goal.

In the present study, we generated a spatially resolved large-scale single-cell open chromatin and transcriptomic map of the adult primate cortex. We applied snATAC-seq and snRNA-seq to profile 358,237 single cells from three major cortical regions; prefrontal cortex, motor cortex and primary visual cortex, and performed the Stereo-seq on these cortical regions to obtain more than 30,000 spots (bins) per region. Relative to single gene or more qualitative approaches, multi-omics data applied in integrated analyses frameworks can increase both resolution and confidence of cell type annotations. To demonstrate the robustness of our data resource, we performed co-embedding of single-cell ATAC-seq and snRNA-seq data that confirmed the high consistency of our cell type assignments and linked the cell type-specific expression profiles to their corresponding regulatory programs.

To map cell types and their molecular properties in their spatial contexts and to investigate cortical area heterogeneity *in situ* we transferred the snRNA-seq data to Stereo-seq data. To our knowledge, this is the first study that integrates transcriptomic with spatial transcriptomics to profile the gene expression map on primate cortical areas. Through this approach, we were able to define the area specific expression and gradient gene expression pattern of excitatory neurons *in situ*. Thus, our combined epigenetic and transcriptomic analysis, spatially resolved with Stereo-seq profiling, synergized to reveal hidden aspects of spatial organization and cellular heterogeneity in cerebral cortex.

Dysfunctional myelination is a feature in several neurodegenerative diseases and neurodevelopmental disorders, including multiple sclerosis^76^. Our data resource allowed us to perform an integrative analysis on single cell transcriptomic and epigenetic profiling of OPC and OLIs, resolving a parallel dynamic gene expression landscape, open chromatin states and transcription factor enrichments related to myelination. By applying pseudotime analysis, we were able to map how specific gene expression changes, chromatin states and regulatory circuitries influence cell fate decisions throughout lineage maturation, define roles of master regulators, and identify potential targets for demyelination diseases. In addition, comparative analysis of the OLI lineage trajectories revealed the shared and divergent gene activation and TFs regulations between macaque and human.

A single-cell macaque genomics data resource can be applied directly in pre-clinical studies using NHP models of neuropsychiatric and neurological diseases. Here, we used our cell-type specific epigenetic data and the spatially resolved Stereo-seq data to predict the risk enrichment of neurological and neuropsychiatric disorders. Notably, mapping heritability loci with chromatin accessibility data, and mapping cell type perturbation with spatial transcriptomic data can help us understand whether or not aspects of disease biology is recapitulated in different cell types or brain regions. For example, in the PFC region, we found a highly overlapping cell type enrichment for genetic risk and disease-perturbation genes for SCZ, but not for ASD. Even for SCZ, the overlapping cell type enrichment for these two features varied between PFC, M1 and V1 such that the overlap of cell types in PFC and M1 was larger than it was in V1. Thus, our multi-omics macaque cortical cell data provide a valuable resource that can readily be applied to map human heritable traits and compared with disease model data.

Future studies on tissues from cortical and sub-cortical areas from developmental and adult stages, subjected to integrative analysis of single cell transcriptomic, epigenetic, proteomics and spatial genomics promise to uncover systemic molecular mechanisms of the primate brain in health and disease. With the first dataset at this high resolution, we identified regulatory elements and spatial expression profiles that shape primate cortical organization and harnessed valuable insights with relevance to human disease.

## Extended Data Figures

**Extended Data Figure 1.**
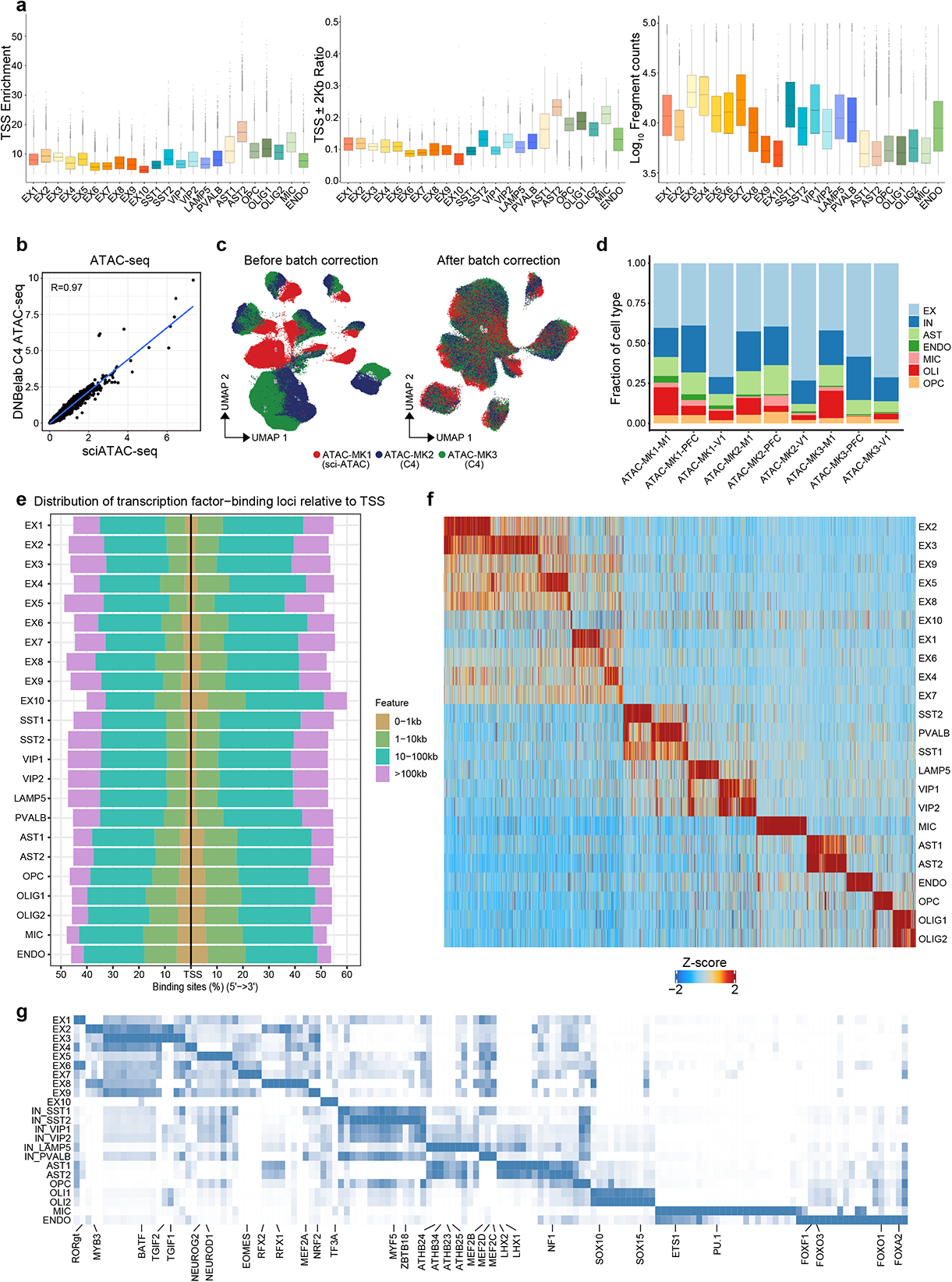
Quality assessment of snATAC-seq data. **a,** Box plot of TSS enrichment: average accessibility of the TSS +/- 50 bp region/ average accessibility of the TSS flanking positions (+/- 1900 – 2000 bp) (left), TSS ± 2 kb ratio: the proportion of peaks within 2 kb from the gene TSS site (middle) and fragment counts (right) of snATAC-seq cells in each snATAC-seq cluster. **b,** Correlation between sciATAC-seq data and DNBelab C4 snATAC-seq data, R value was calculated by Pearson correlation. **c,** UMAP projections of all primary snATAC-seq cells before (left) and after (right) batch correction colored by individual donor (the library preparation methods are indicated). **d,** Proportion of each major cell type in each cortical region of each individual donor from snATAC-seq cells. **e,** Stacked columns indicate the genomic distance distribution of differential accessible *cis*-elements (ATAC-seq peaks) in each snATAC-seq cluster. **f,** Heatmap showing the differentially accessible *cis*-regulatory elements in each snATAC-seq cluster. **g,** Heatmap showing the top 30 transcription factor (TF) binding motifs enriched at differential accessible peaks of each snATAC-seq cluster (Fig. 1b) by HOMER.

**Extended Data Figure 2.**
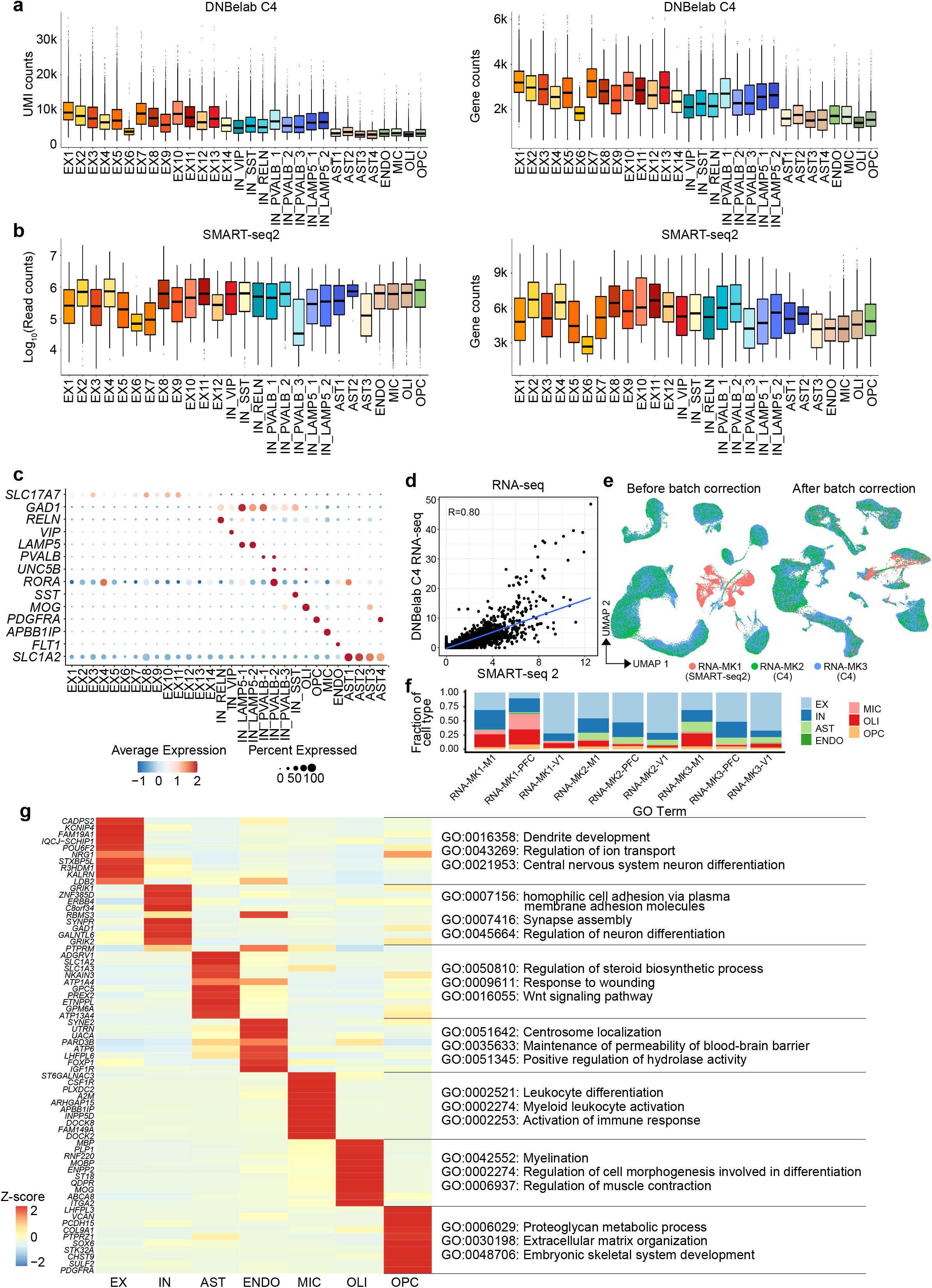
Quality assessment of snRNA-seq data. **a,** Box plot of unique molecular identifiers counts (UMI) (left) and detected gene number (right) of DNBelab-C4 cells in each snRNA-seq cluster. **b,** Box plot of reads count (left) and detected gene number (right) of Smart-seq2 cells in each snRNA-seq cluster. **c,** Dot plots illustrating marker gene expression in snRNA-seq cell clusters. Size of each dot represents the percentage of cells expressing the marker. **d,** Correlation between SMART-seq2 data and DNBelab C4 snRNA-seq data, R value was calculated by Pearson correlation. **e,** UMAP projections of all primary snRNA-seq cells before (left) and after (right) batch correction colored by individual donor (the library preparation methods are indicated). **f,** Proportion of each major cell type in each cortical region of each individual donor from snRNA-seq cells. **g,** Heatmap displaying the differentially expressed genes of major cell classes in snRNA-seq data. Specific genes related to each cell type are highlighted with enriched gene ontology terms.

**Extended Data Figure 3.**
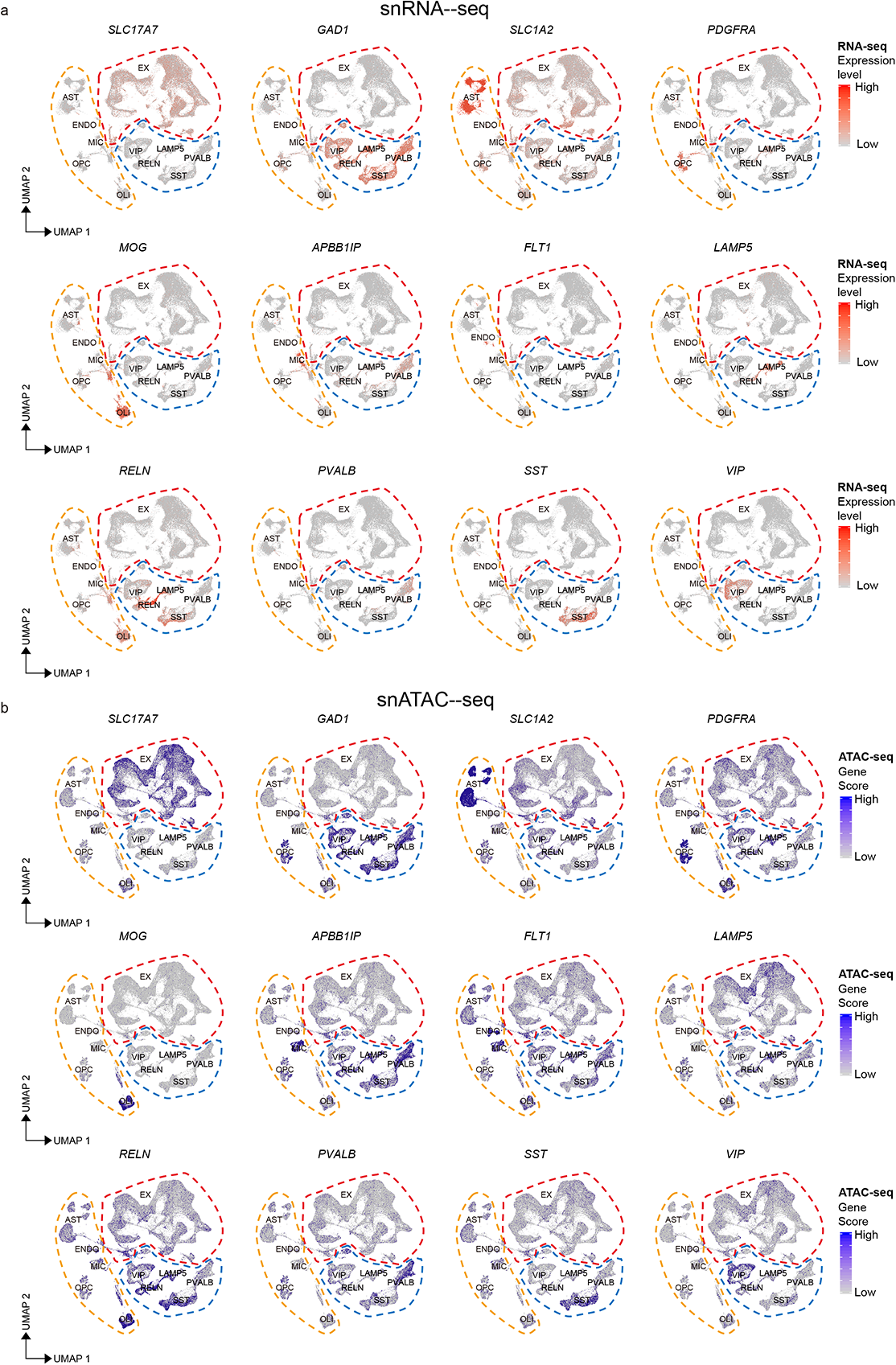
Congruent cell type identifications in snRNA-seq cells and snATAC-seq cells. **a,** Expression levels of cell type-specific marker genes in snRNA-seq cells visualized by UMAP. **b,** Gene activity score of cell type-specific marker genes in snATAC-seq cells visualized by UMAP.

**Extended Data Figure 4.**
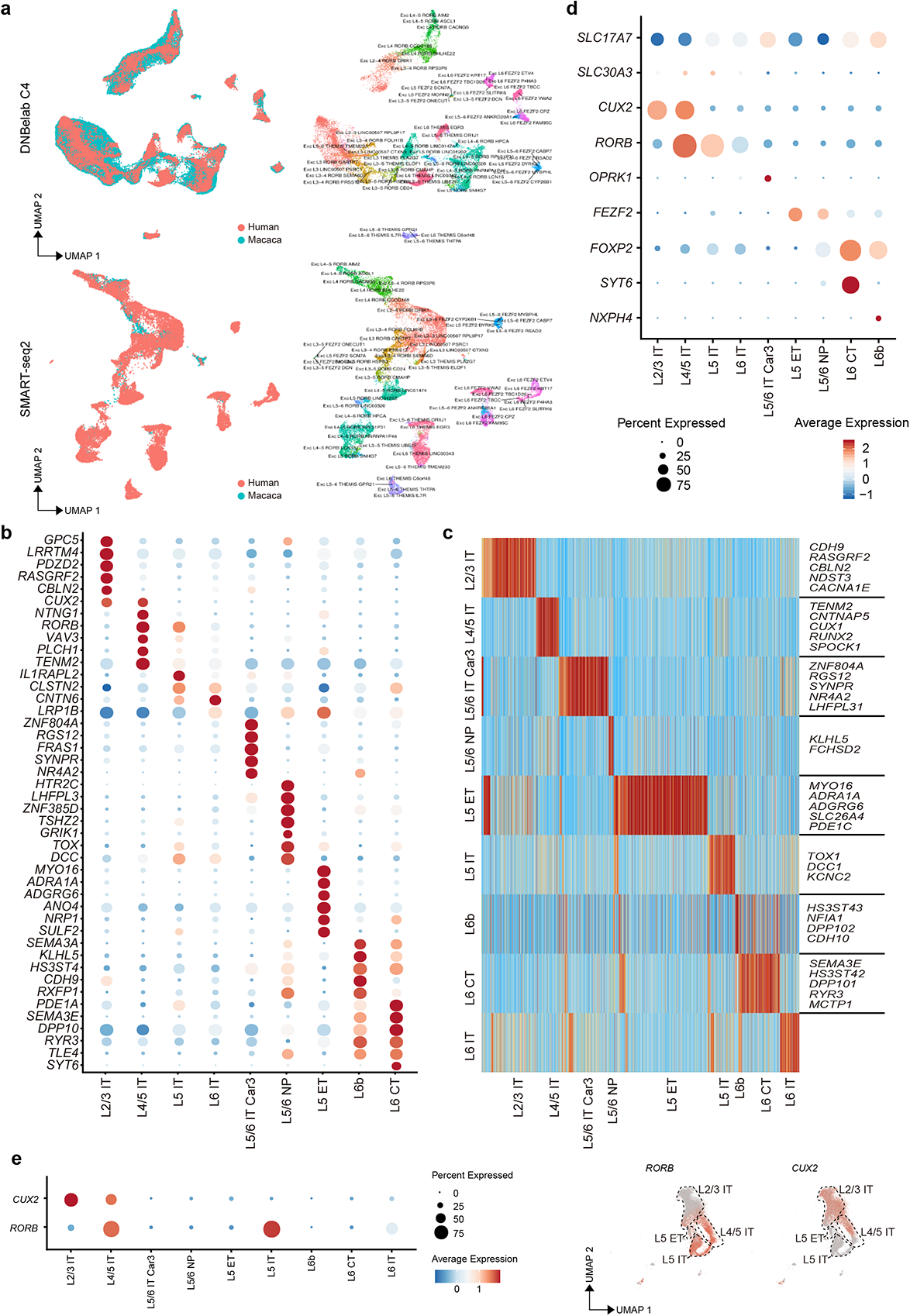
Chromatin accessibility and gene expression of excitatory neurons subtypes. **a,** Integration of DNBelab C4 (top) and Smart-seq2 (bottom) snRNA-seq data of excitatory neurons from macaque with single-nucleus transcriptomes of human cortex (left), and cluster labels of human excitatory neurons were indicated (right). **b,** Dot plots showing expression of marker genes across macaque excitatory neuron cell subtypes of snRNA-seq. **c,** Heatmap showing the differentially accessible peaks enriched across macaque excitatory neuron subtypes of snATAC-seq. **d,** Expression of marker genes for mouse excitatory cell classes in macaque excitatory neuron subtypes. **e,** Dot plots (left) and feature plots (right) showing layer marker of L2-4 (CUX2) and L3-5 (RORB) are co-expressed in L4/5 IT of M1.

**Extended Data Figure 5.**
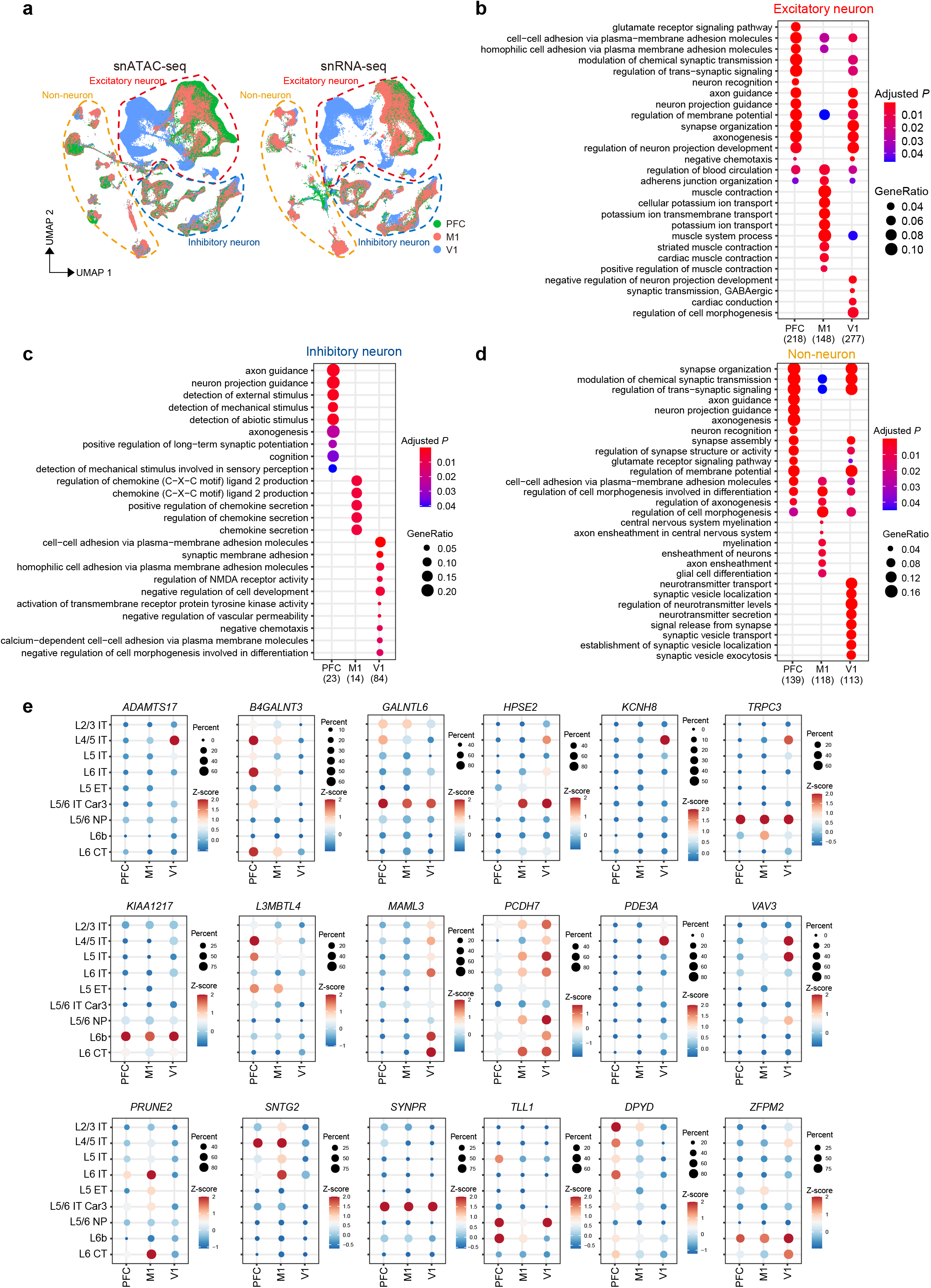
Areal differences within excitatory neurons. **a,** UMAP plot of snATAC-seq cells (left) and snRNA-seq cells (right) colored by cortical areas. **b,** Gene ontology terms enriched among differentially expressed genes in PFC, M1 and V1 of excitatory neuron subtypes. **c,** Gene ontology terms enriched among differentially expressed genes in PFC, M1 and V1 of inhibitory neuron subtypes. **d,** Gene ontology terms enriched among differentially expressed genes in PFC, M1 and V1 of non-neuronal cell types. **e,** Dot plots of selected differentially expressed genes in PFC, M1 and V1 of excitatory neuron subtypes.

**Extended Data Figure 6.**
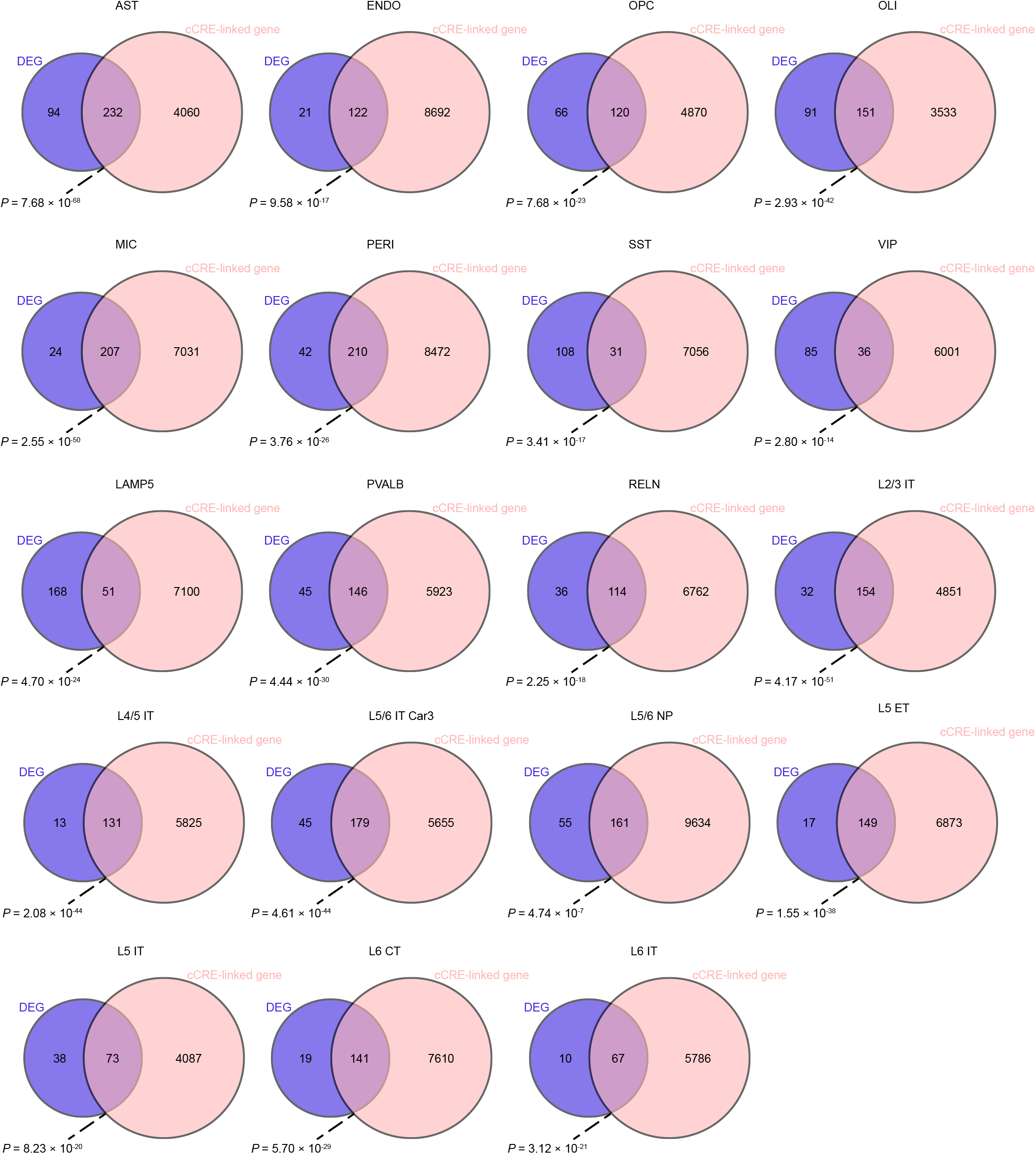
Linking *cis*-regulatory elements to cell type specific genes. Venn diagrams showing the overlaps between CRE-targeted genes and differentially expressed genes (DEGs) in that cell type/subtype.

**Extended Data Figure 7.**
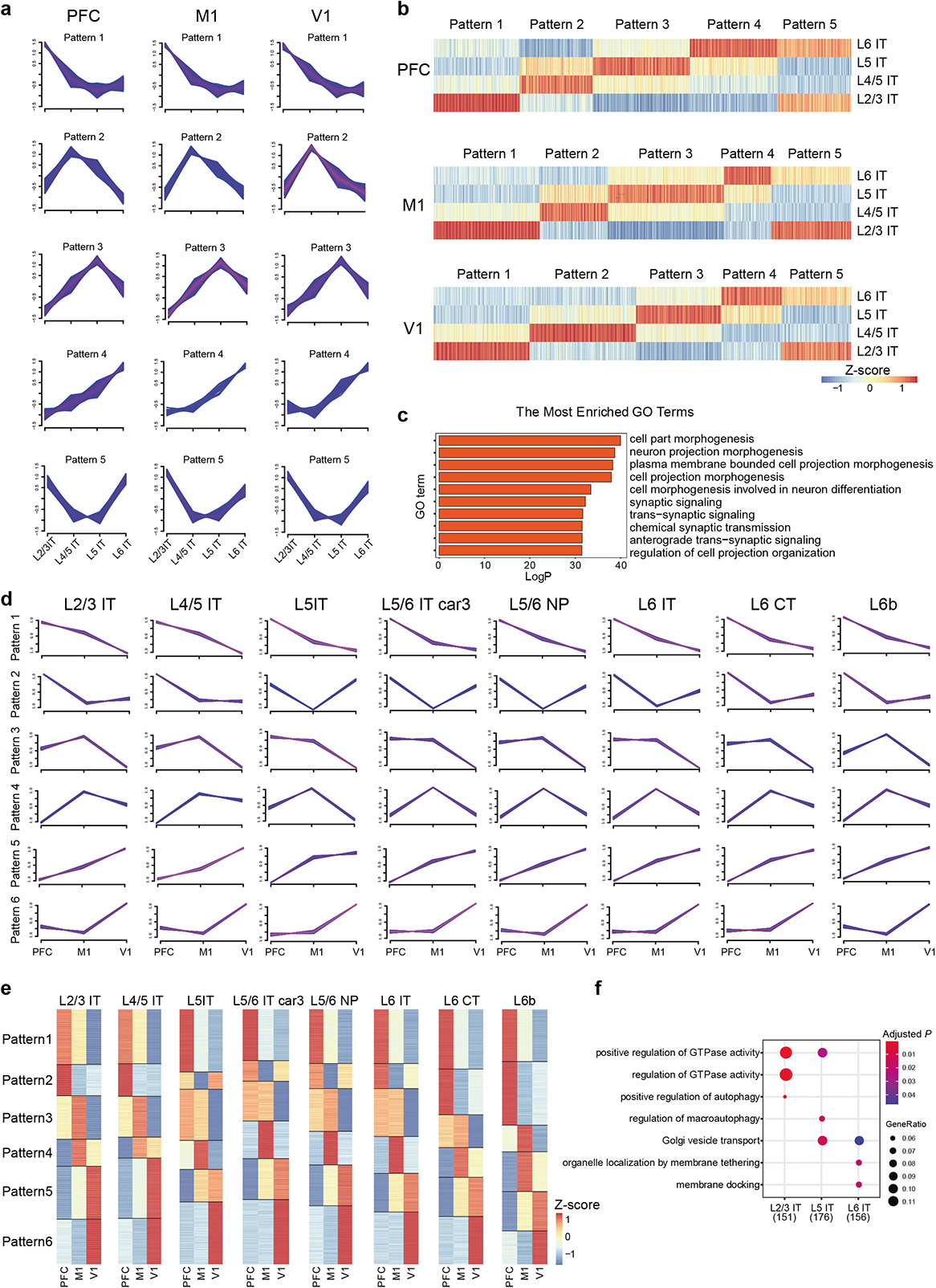
Gradient gene expression pattern of excitatory neurons. **a,** Gradient genes expression patterns across L2/3 IT, L4/5 IT, L5 IT and L6 IT by Mfuzz in PFC, M1 and V1 (probability of genes matching the pattern > 0.5). **b,** Heatmap showing the genes from a) and filtered by maximum expression > 1. **c,** Gene ontology terms enriched among genes with distinct expression pattern between PFC, M1 and V1. **d,** Gradient gene expression patterns across PFC, M1 and V1 by Mfuzz in excitatory neuron subtypes (probability of genes matching the pattern > 0.5). **e,** Heatmap showing the genes from d) and filtered by maximum expression > 1. **f,** Gene ontology terms enriched among genes in pattern 1 of L2/3 IT, L5 IT and L6 IT types.

**Extended Data Figure 8.**
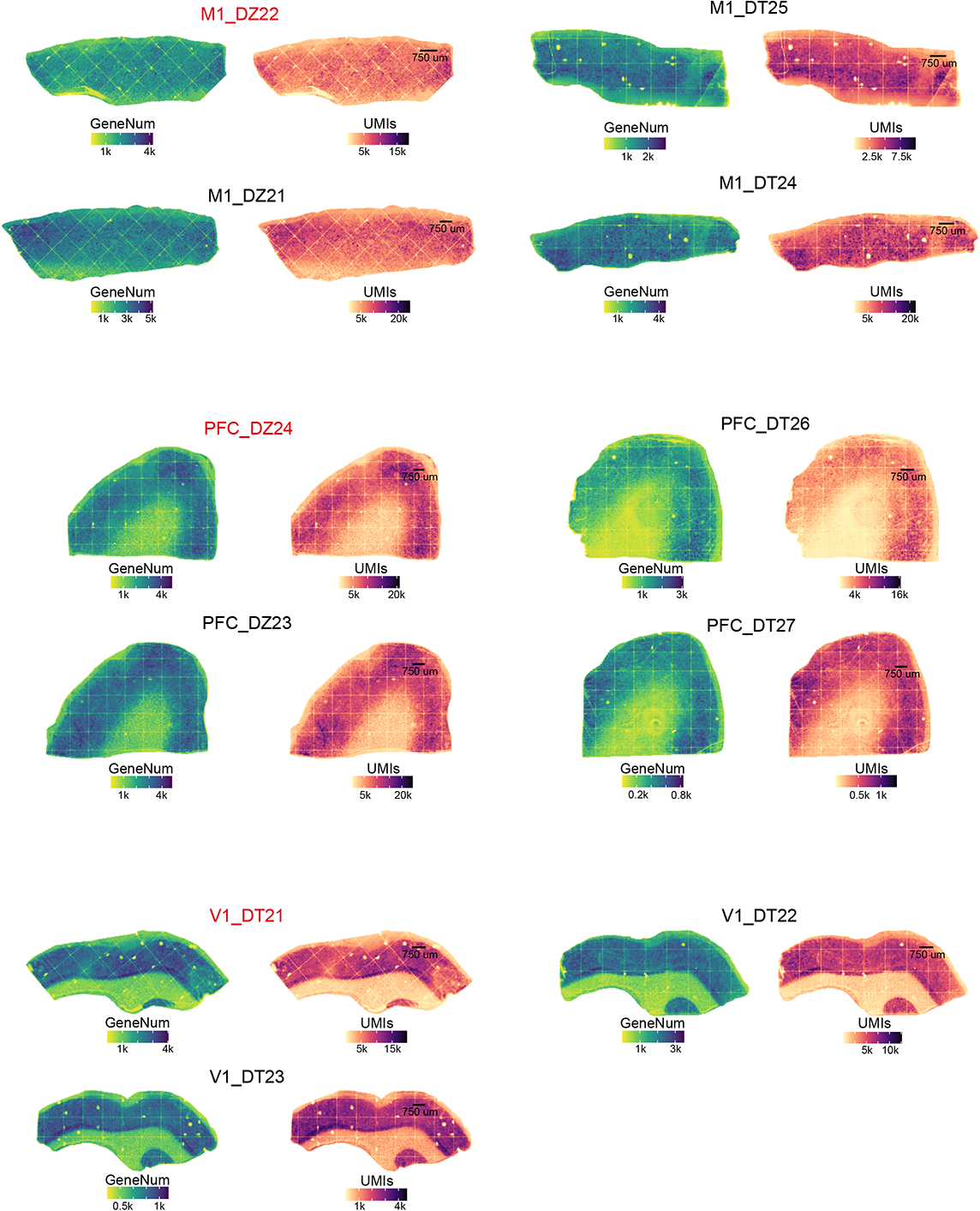
Detected gene number and UMI counts of Stereo-seq sections. Spatial distribution of detected genes number and UMI counts per 37.5 μm bin in Stereo-seq sections from PFC, M1 and V1. Sections used for further analysis were red color coded.

**Extended Data Figure 9.**
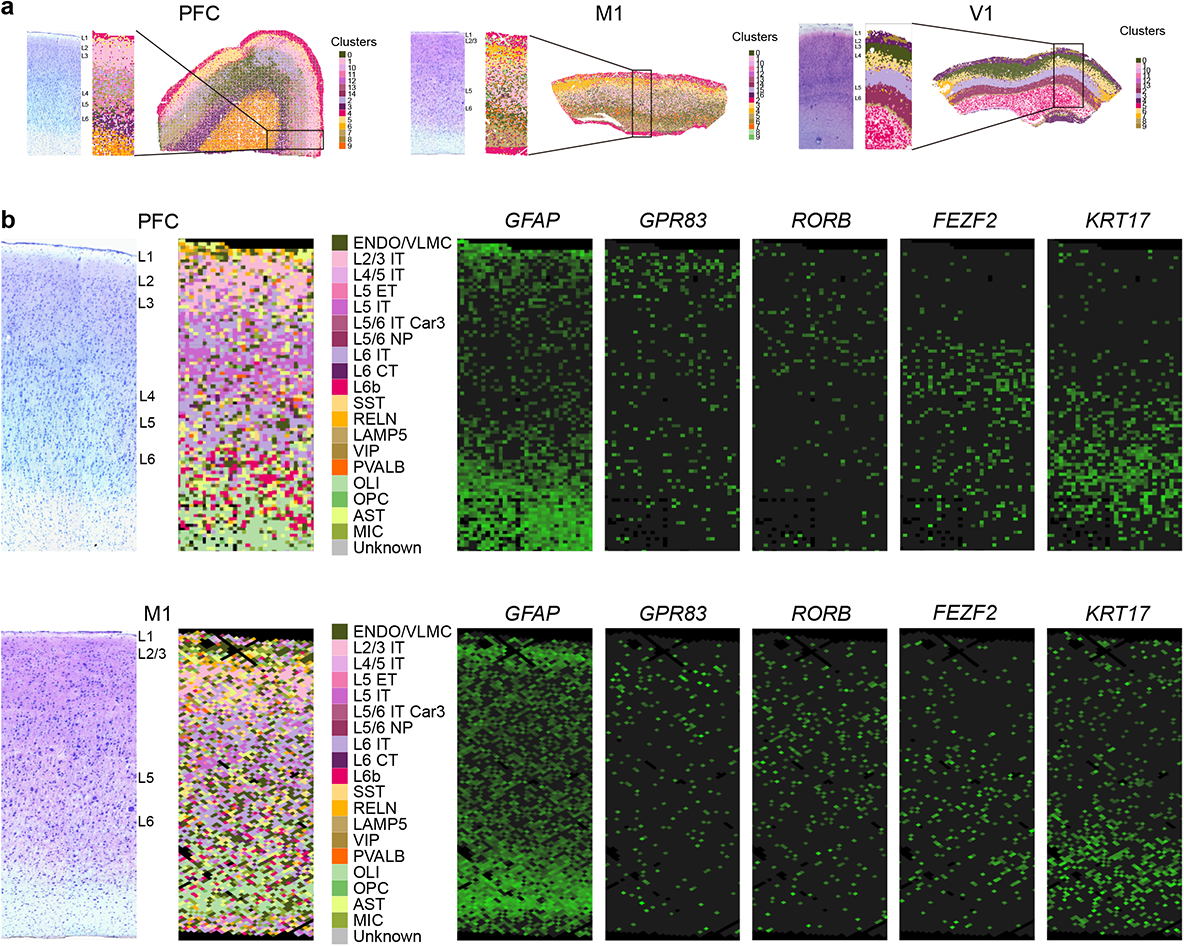
Unsupervised clustering and cell type annotation of Stereo-seq sections. **a,** Nissl staining of adjacent section to stereo-seq section and unsupervised clustering of Stereo-seq spots (37.5 μm bins) of PFC, M1 and V1. **b,** Nissl staining of adjacent section to stereo-seq section of PFC (top left) or M1 (bottom left), cell type/subtypes annotation of stereo-seq spots (37.5 μm bins) by snRNA-seq data (middle), and layer enrichment of known layer-marker genes in the same stereo-seq section (right).

**Extended Data Figure 10.**
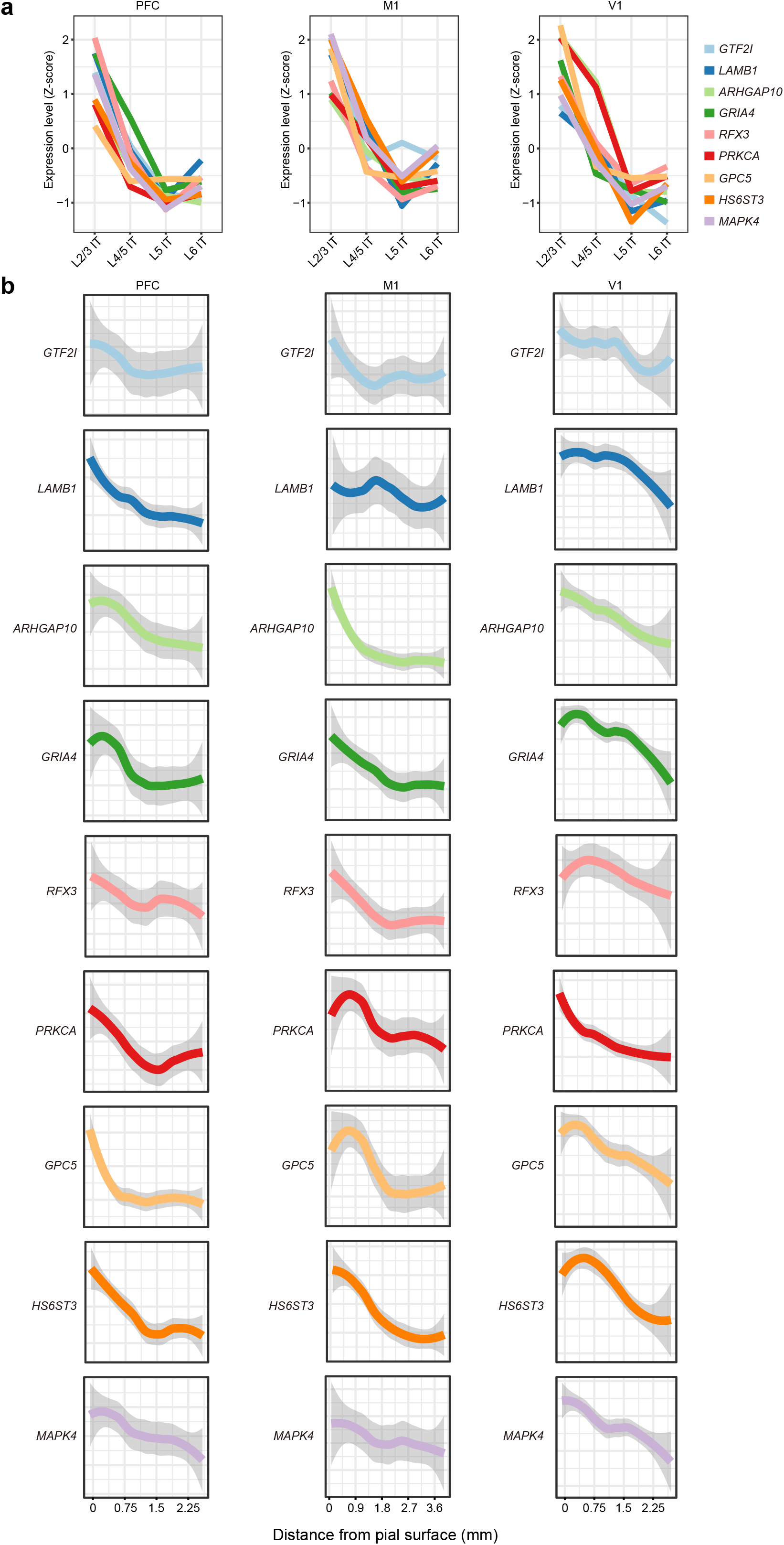
Gradient expressed genes in excitatory neuron of Stereo-seq sections. **a,** Genes showed congruent gradient expression pattern (pattern 1 in Extended Data Fig. 7a) between PFC, M1 and V1 of snRNA-seq cells. **b,** Gradient expression of genes from **a)** in excitatory neuron of Stereo-seq sections from PFC, M1 and V1.

**Extended Data Figure 11.**
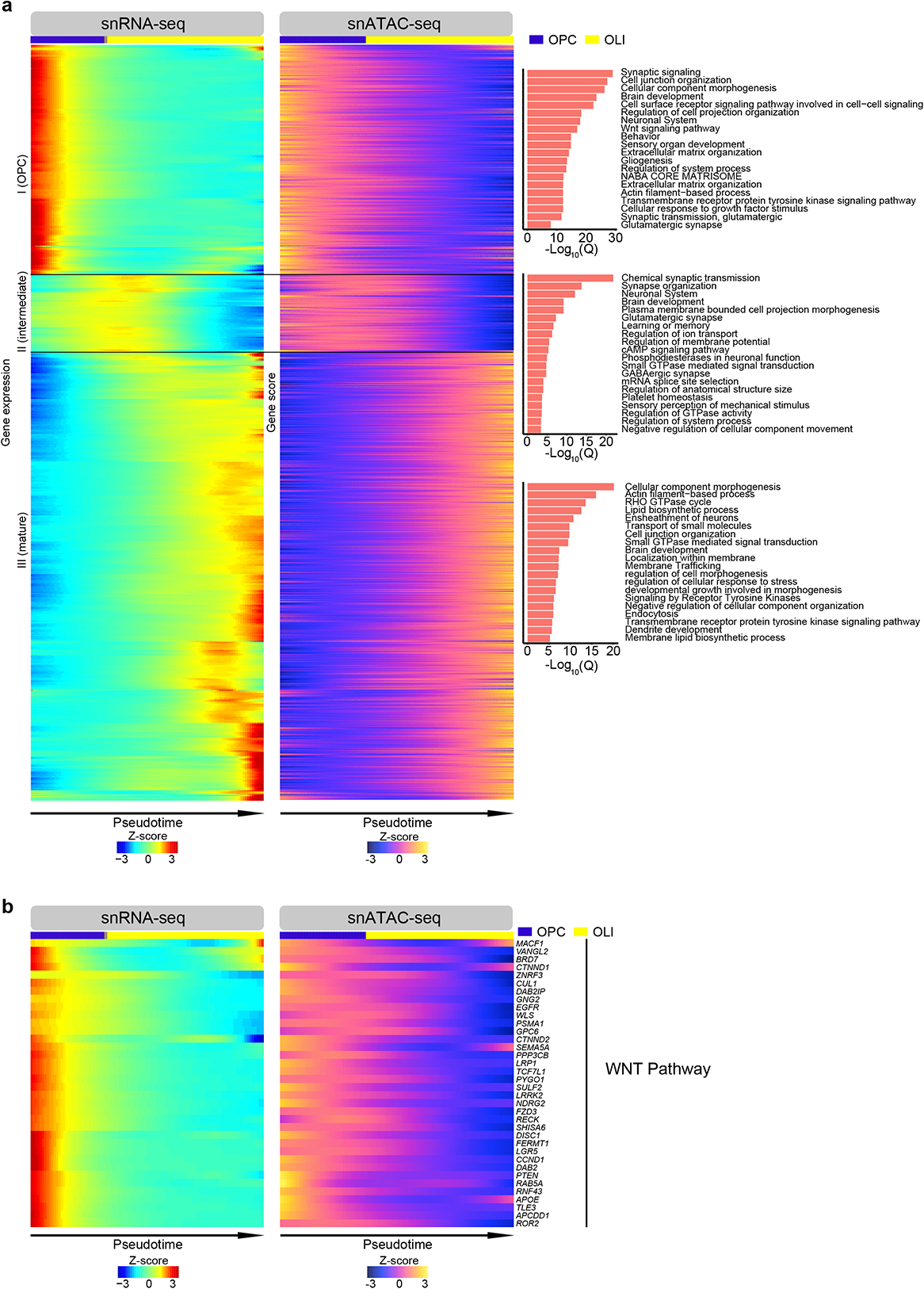
Well-concordance gene expression-gene activity score pairs along macaque OLI lineage trajectory. **a,** Well-concordance gene expression (left)-gene activity scores (middle) pairs and ontologies terms (right) along macaque OLI pseudotime trajectory (Spearman’s correlation coefficient r_s_ > 0.2, P < 0.01). **b,** Well-concordance gene expression (left)-gene activity score (right) pairs of genes involved in Wnt signaling pathway along macaque OLI pseudotime trajectory.

**Extended Data Figure 12.**
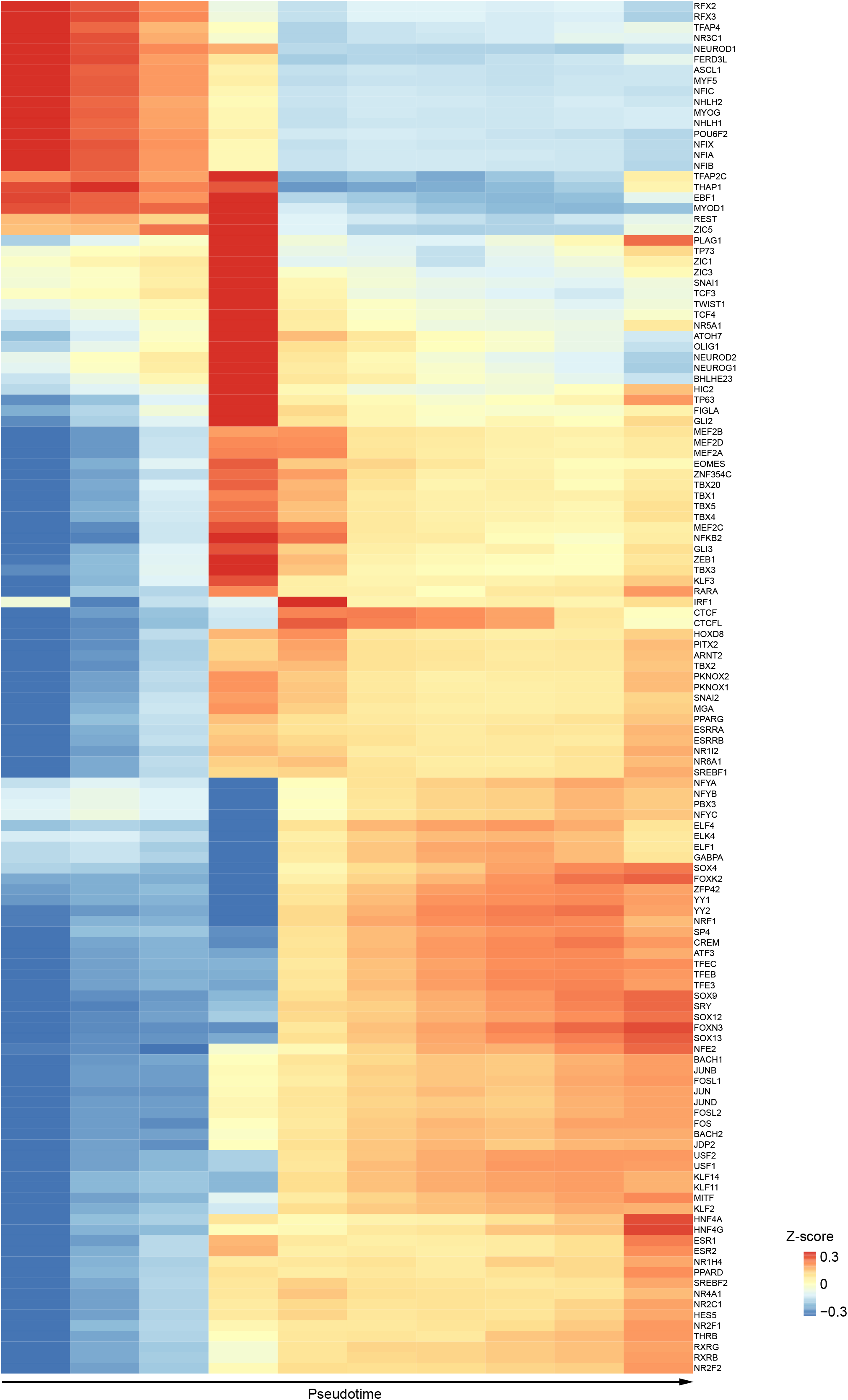
TF motif enrichment along macaque OLI lineage trajectory. Heatmap ordering of average TF binding motif bias-corrected deviations for 129 most variable TFs along macaque OLI pseudotime.

**Extended Data Figure 13.**
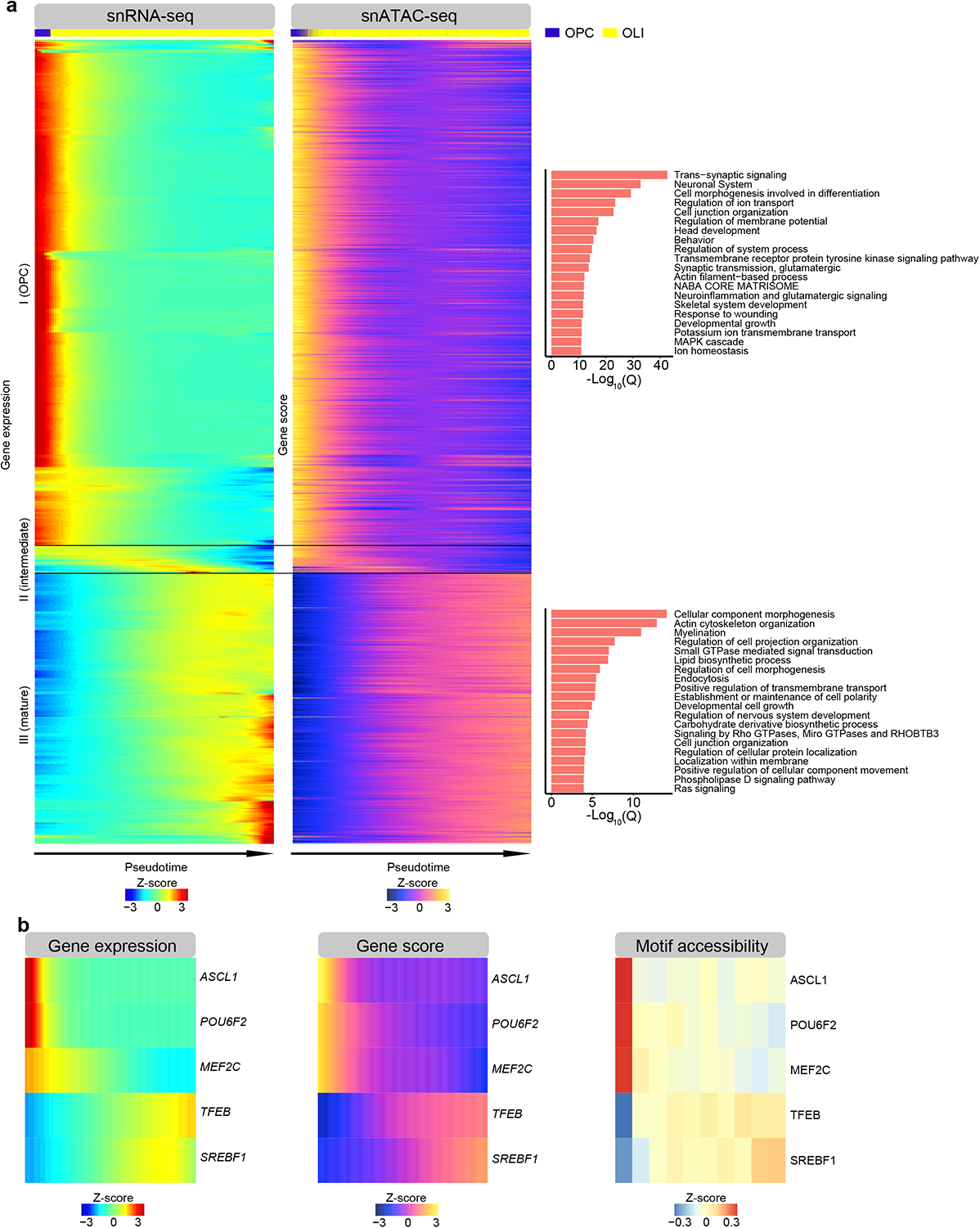
Well-concordance gene expression-gene activity score pairs along human OLI lineage trajectory. **c,** Well-concordance gene expression (left)-gene activity scores (middle) pairs and ontologies terms (right) along human OLI pseudotime trajectory (Spearman’s correlation coefficient rs > 0.2, P < 0.01). **d,** Gene expression (left), gene activity score (middle) and motif enrichment (right) of 5 TFs across human OLI pseudotime trajectory.

## Supplementary Tables

**Supplementary Table 1. Sample annotation and quality information of snRNA-seq and snATAC-seq data**

**Supplementary Table 2. Top 30 transcription factor (TF) motifs enrichment in each snATAC-seq cluster**

**Supplementary Table 3. Differentially expressed genes among snRNA-seq clusters.**

**Supplementary Table 4. Differentially expressed genes and differentially accessible peaks of layer/projection-defined 9 EX subtypes.**

**Supplementary Table 5. Differentially expressed genes between PFC, M1 and V1 in each cortical cell type/subtype.**

**Supplementary Table 6. Differentially accessible peaks between PFC, M1 and V1 in each cortical cell type/subtype.**

**Supplementary Table 7. Differentially expressed genes between PFC and M1 in excitatory neuron subtypes.**

**Supplementary Table 8. Gradient gene expression patterns across L2/3 IT, L4/5 IT, L5 IT and L6 IT by Mfuzz in PFC, M1 and V1.**

**Supplementary Table 9. Gradient gene expression patterns across PFC, M1 and V1 by Mfuzz in excitatory neuron subtypes.**

**Supplementary Table 10. Differentially expressed genes between PFC, M1 and V1 in excitatory neuron subtypes of snRNA-seq also area-enriched in Stereo-seq sections.**

**Supplementary Table 11. Well-concordance set of genes in expression-activity score pairs along oligodendrocyte pseudotime trajectory in macaque monkey and human.**

**Supplementary Table 12. Summary of GWAS studies of psychiatric disorders and neurobehavioral traits used for disease risk mapping.**

## Methods

### Ethics Statement

All relevant procedures involving animals were approved in advance by the Institutional Animal Care and Use Committee of Yunnan Key Laboratory of Primate Biomedical Research.

### Tissue processing and Nuclei isolation

Tissues were sampled from three female 72-month-old cynomolgus monkeys (Macaca fasicularis) and immediately frozen in liquid nitrogen. Single-nucleus preparations were performed as previous description^77^, frozen monkey brain tissue pieces were placed in 1 ml homogenization buffer (pre-chilled) in 1 ml Dounce homogenizer (TIANDZ). Tissue was homogenized by 10 strokes of the loose pestle and 10 strokes of the tight pestle, Dounce homogenizer was submerged in ice during grinding. 2 ml homogenization buffer was added to the Dounce homogenizer then the homogenate was filtered through 40μm cell strainer (Miltenyi Biotech) into 15 ml conical tube and centrifuged at 900 g for 10 mins to pellet nuclei.

### snATAC-seq library preparation and sequencing

Sci-ATAC-seq was performed as described previously with modifications^78^. Nuclei were strained in 40 µm strainer and centrifuged for 5 min at 500 g. The nuclei were resuspended in cold PBS (1% BSA) and counted using hemocytometer. Nuclei were adjusted concentration to 360/μL. For transposition, added 7 μL cell suspension (around 2500 cell), 2 μL 5xTAG buffer and 1 μL unique barcoded Tn5 transposome into each well of 96-well plate, mixed gently and had a short spin ^79^. The plate was incubated at 55 for 60 min with shaking (300 rpm). To quench the reaction, 10 μL of 40 mM EDTA was added to each well and gently mixed, then the plate was incubated at room temperature for 5 min. After reaction, 5 μL sorting buffer (5% BSA; 5 mM EDTA) was applied to each well, mixed well and pooled into one tube. The suspension was filtered through 40 µm strainer. Then one drop of DAPI (4′, 6-diamidino-2-phenylindole, ThermoFisher Scientific) was added to the suspension and 25 nuclei were sorted by Aria II (BD) into 96-well plate containing 7 μL buffer EB, shortly spun down. Next, 1 μL 10% SDS was added to each well, mixed well and incubated at 55c for 7 min with shaking (500 rpm) to lyse the nuclei. After the reaction, 1 μL 10% Triton-X was added to each well, spun down and incubated at room temperature for 5 min. For amplification, we added 1μL unique barcoded N5&N7 (0.5 µM final concentration) and 10 μL NEBNext High-Fidelity 2x PCR Master Mix (NEB). PCR cycling conditions were as follow: 72 °C 5 min, 98 °C 30 s, (98 °C 10 s, 63 °C 30 s, 72 °C 30 s) × 11, held at 4 °C. After that, we pooled each 96-well plate into tubes, and added 5 volume PB including pH-indicator (Qiagen) and 200 μL sodium-acetate (3M, pH = 5.2), and reversed blending. Next, we used 4 columns to purify product following the MinElute PCR Purification Kit manual (Qiagen). DNA from each column was eluate using 25 μL EB buffer, pooled the 4 elution, and added EB buffer to 100 μL. To filtrate the fragment, we used the Ampure XP Bead by 0.5x & 0.7x. First added 50 μL XP beads, after incubation, collected supernatant and then added 70 μL XP beads. Finally, we used 100 μL EB buffer to elute the DNA. We quantified the libraries by Qubit fluorimeter (Life technologies) and detected fragment size using 2100 High Sensitivity (Agilent). To sequence, each library we used 330 ng for cyclizing, and then used 8 ng to make DNB. Each library was loaded into 2 lanes using BGISEQ500. The sequencing primer used as: Tn5 primer1, Tn5 primer2, SCIMDA primer. The read lengths: PE 100, including 4 indexes. Index 1 and index 2 was represented Tn5 barcode, index 3 and index 4 represented the PCR barcode. There are common bases between Tn5 barcode and PCR index, so we used cold reaction of BGISEQ500 to get out of the imbalance.

Drop-seq based DNBelab C4 ATAC-seq was performed using DNBelab C Series Single-Cell ATAC Library Prep Set (MGI, #1000021878) with the procedure of step transposed, droplet encapsulation, pre-amplification, emulsion breakage, then the captured beads were collected. After DNA amplification and purification, the snATAC library is already to sequence. DNA nanoballs were loaded into the chips and sequenced on the DIPSEQ T1 sequencer at China National Gene Bank (CNGB).

### snATAC-seq data processing and quality control

We used an in-house pipeline to process the sci-ATAC-seq data. Briefly, we extracted the segments of barcode from the 4 constituent parts (read 1: 1-10, 32-41; read 2: 1-10, 38-47), and retained the reads with all segments of barcode are fully matched. Then we mapped the reads to *macaca fascicularis* genome downloaded from NCBI using bowtie2 ^80^ with ‘-X 2000 --mm --local’ as options. Reads with mapping quality less than 10 and reads mapped to the mitochondria or genome scaffold (chrAQ*, chrU*, chr*_random*, and chrK*) were filtered out. Then, PCR duplicates were removed according to the cell barcode and mapping loci by custom python script. The retained fragments of each library were used for further analysis.

We used an open-source pipeline (https://github.com/shiquan/PISA) to analyze the DNBelab C4-based snATAC-seq data. First, the raw reads were filtered and demultiplexed using PISA with 2 mismatches allowed in barcode. Then the retained reads were aligned to the *macaca fascicularis* genome using BWA-MEM () with the default parameters. Reads with mapping quality less than 10 and reads mapped to the mitochondria or genome scaffold (chrAQ*, chrU*, chr*_random*, and chrK*) were filtered out and PCR duplicates were removed according to the cell barcode and mapping loci. The retained fragments of each library were used for further analysis.

### snATAC-seq data clustering and analysis

We applied ArchR (https://github.com/GreenleafLab/ArchR) to identify the cell populations within all data. First, cells with fragments lower than 3000 and TSS enrichment lower than 4 will been filtered, then doublet score were calculated via the addDoubletScores function and filtered the doublets by the filterDoublets function with parameter: filterRatio = 2. Before defined cell clusters, we first created 500 bp tiled matrix of the genome and used the addIterativeLSI function to reduce to 30-dimension base on the top 15000 most accessible site across all cells. Then used the AddHarmony to correct the batch base on donors and different cortical regions and identified the clusters using Seurat’s SNN graph clustering with a default resolution of 0.8. Then peaks of each cluster were calculates using the addReproduciblePeakSet function with macs2, and differential peaks of each cluster were identified by getMarkerFeatures function.

### snRNA-seq library preparation and sequencing

For Smart-seq2 processing, nuclei were resuspended in 500 μL blocking buffer containing 1X PBS (GIBCO), 2% filtered sterilized BSA (SIGMA), and 0.2 U/μL RNasin Plus by pipetting up and down gently on ice. Nuclei were transferred to 1.5 ml tube then split up into three tubes: experimental, control-isotype and control-DAPI. For single-nuclei isolation by flow cytometry, 250 μL DAPI (0.1 μg/ml) was added into the experimental tube to make the final volume as 500 μl, while nothing added to the negative control tube. We also sorted NeuN^+^ nuclei from M1 region. For this purpose, rabbit anti-NeuN (Abcam, final dilution of 1:500) was added to experimental tube, rabbit IgG-monoclonal-isotype control (Abcam, final dilution of 1:500) was added to control-isotype tube. After 30 min incubation at 4 °C, samples were then centrifuged for 5 min at 400 g to pellet nuclei and pellets were resuspended in 250 μL blocking buffer (including 50 μL leaving buffer). 250 μL DAPI (0.1 μg/ml) was added to nuclei to make the final volume for each tube as 500 μL. Stained nuclei were filtered through 40 μm filter before FACS (BD FACSAria II instrument). DAPI or DAPI^+^ NeuN^+^ nuclei in the experimental tube were sorted at a speed of 5-7 and the single nucleus was sorted into every single well of 384 plate filled with 1.2 μL lysis buffer (10% Triton X-100 0.0125 μl, 40 U/μl RNase Inhibitor 0.0625 μl, 10 M Oligo-dT Primer 0.02 μL, 10 mM dNTP Mix 0.45 μL, 5× SuperScriptII First-Strand Buffer 0.25 μL, nuclease-free water 0.305 μL and ERCC spike-in 0.1 μL) in advance. Sorted nuclei in 384-well plates were briefly centrifuged and stored at −80 °C for further analysis. Sorted nuclei transcriptome amplifications were prepared by a modified Smart-seq2 protocol 81. After nuclei lysis, 1.2 μL reverse transcription mixed solution was added into each well to complete the reverse transcription reactions. Then 1.8 μLPCR reaction buffer was added into every well to complete amplification. At last, the amplified cDNA products of each single nucleus were quantified by Agilent Bioanalyzer 2100. For those single nucleus samples with high quality after amplification, the products were extracted by an automatic extractor from the 384-well plate to 96-well plate then purified by MGIEASY DNA Purification Magnetic Bead Kit (MGI) for the library construction. The libraries were prepared by MGIEASY RNA Library Preparation Kit (MGI) and each single nucleus sample was barcoded. Finally, the libraries were cyclized into ssDNA libraries by MGIEASY Cyclization Kit (MGI). All the single nucleus samples were sequenced on the BGISEQ500 sequencer with 100-bp pair-end reads.

For drop-seq based DNBelab C4 RNA-seq processing, snRNA-seq libraries were prepared using DNBelab C Series Single-Cell Library Prep Set (MGI, #1000021082). The procedure was utilized as previously described^27^. Briefly, we followed the manufacturer’s protocol for droplet generation using the single nucleus suspensions, emulsion breakage, beads collection, reverse transcription and cDNA amplification to generate barcoded libraries. The resulting libraries were sequenced using DIPSEQ T1 sequencers at the China National GeneBank (Shenzhen, China). The read length followed: read 1, 30 bp, including 10 bp cell barcode 1, 10 bp cell barcode 2 and 10 bp UMI, read 2, 100 bp for transcript and 10 bp for sample index.

### snRNA-seq data processing and quality control

Smart-seq2 data were processed as previous described^81^, first, one-hundred-base-pair paired-end reads were pre-processed to remove adapters and filter out reads with low quality using default parameter by Cutadapt (v1.15) ^82^. Next, filtered reads were aligned to the Macaca fascicularis genome (5.0.91) by STAR (v2.5.3) ^83^ using a modified GTF file from ensemble release-91. In order to map the pre-mRNA fragments which may cover both exonic and intronic regions, we create a modified GTF annotation file which only contain transcript regions, and the original annotation rows for exons were all deleted, then we replace the feature type name from ‘transcript’ to ‘exon’. Finally, transcripts per million mapped reads (TPM) were calculated using rsem-calculate-expression (RSEM) ^84^ with default parameters.

We applied PISA (version 0.7) to filter and demultiplex DNBelab C4 RNA-seq raw sequencing reads, then used STAR (version 2.6.1a) to align the reads with the previously mentioned modified GTF file of Macaca_fascicularis_5.0 genome containing introns and exons and sort by sambamba (version 0.7.0). For each library, we sorted the cells from high to low according to the number of UMIs and kept the cells whose UMI number is greater than the UMI of the tenth-ranked cell divided by 10, and then used PISA to generate the UMI count matrix of the cell to the gene. For Smart-seq2 data, cells with unique mapped ratios > 15% and more than 500 genes with transcripts per million mapped reads value >1 and removed the top 5% of cells with the highest number of genes were used for downstream analysis. For DNBelab C4 data, cells with more than 500 genes and removed the top 5% of cells with the highest number of genes were used for downstream analysis. For both Smart-seq2 and DNBelab C4 RNA-seq data, cells with mitochondrial gene expression greater than 1% were filtered, and then all mitochondrial genes were deleted for downstream analysis.

### snRNA-seq data clustering and analysis

Clustering and differential gene expression analysis was performed by Seurat (v 3.1.1)) R toolkit^85^. NormalizeData, FindVariableGenes and ScaleData were performed respectively in each cortical region of brain samples. Genes expressed in less than three cells were filtered out and cells with expressed genes less than 500 were excluded. SCtransform (SCT) was performed to integrate cells from 3 monkeys using the union set of 2000 variable genes in 3 batches. We select the top 20 dimensions (dims = 1-20) for SNN network construction. Then, cell clusters were identified by graph-based clustering approach, louvain algorithm (resolution = 0.5). We used uniform manifold approximation and projection (UMAP) to visualize the distance between cells in 2D space. The edges between two cells were measured by Jaccard distance to construct SNN graph. Then graph-based clustering was performed (times = 50) and modularity was calculated until reaching the maximum. “FindAllMarkers” and “FindMarkers” functions were used for differential gene expression analysis between clusters.

### Annotation of excitatory subtypes by snRNA-seq data of human cortex

The subtypes of excitatory neuron were annotated by co-clustering with human cortical data(https://portal.brain-map.org/atlases-and-data/rnaseq/human-multiple-cortical-areas-smart-seq) excitatory subtypes with R package Seurat (v3.1.1). We integrated the Smart-seq2 and DNBelab C4 data of macaque cortex with human cortical data, respectively by following steps.: Log normalization was performed to integrate cells from each region of human and monkey brain using the union set of 3000 variable genes. We identified anchors using the FindIntegrationAnchors function with a default dimension of 20. We then passed these anchors to the IntegrateData function, following by scaling the integrated data, running PCA, and visualizing the results with UMAP. After finding anchors, we used the TransferDatafunction to classify the macaque cells based on human excitatory subtypes. Cells acquired the label of L2-3 (IT) and L3 (IT) were defined as L2/3 IT type neuron, cells acquired label of L3-4 (IT), L3-5 (IT), and L4 (IT) were defined as L4/5 IT type neuron, cells acquired labels of L4-5 (IT), L4-6 (IT) and L5 (IT) were defined as L5 IT type neuron, cells acquired labels of L5-6 (IT) and L6 (IT) were defined as L6 IT type neuron. Cells acquired labels of L5/6 IT Car3, L5-6 NP, and L6b were defined as the corresponding identities. Cells acquired labels of L3-5 ET and L5 ET were defined as L5 ET type neuron, and cells acquired labels of L5-6 CT and L6 CT were defined as L6 CT type neuron.

### Co-embedding of snRNA-seq cells with snATAC-seq cells

We performed co-clustering of snATAC-seq data and snRNA-seq data with R package Seurat (v3.1.1) ^29, 86^. For snATAC-seq data, we firstly converted the peak by cells matrix to gene activity score. Then, we used RunLSI function with a default dimension of 50 to reduce the dimension. The function of LSI is computed by the term frequency-inverse document frequency (TF-IDF) transformation. We had pre-processed and clustered a snRNA-seq dataset using default parameters as described in “snRNA-seq data clustering” session. Next, we computed anchors between the snATAC-seq dataset and the snRNA-seq dataset by FindTransferAnchors function; and used these anchors to transfer the labels from snRNA-seq data to snATAC-seq cells. Then we used predictions and confidence scores for each snATAC-seq cell correspond to snRNA-seq cell to transfer the cluster IDs that defined in “Annotation of excitatory subtypes by scRNA-seq data of human cortex” session. Next, we created snRNA-seq matrix by transfer cell type labels anchors with TransferData function. We only kept cells with the same cell type definition defined by differential gene activity score and transferred label in ATAC-seq. Finally, we merged measured and imputed snRNA-seq data and snATAC-seq data and run a standard UMAP analysis by reduce dimensionality to 50 to visualize all the cells together.

### Analysis of peak to gene correlations across all cell types

We integrate ATAC-seq data of each cell type/subtype with the RNA-seq data of corresponding cell type/subtype through addGeneIntegrationMatrix function of ArchR, and then use the peak accessibility of ATAC-seq data and the gene expression of RNA-seq data to establish a correlation through addPeak2GeneLinks function of ArchR in each cell type/subtype. To retain reliable correlations, we select correlations for downstream analysis by using correlation coefficients greater than 0.45 and FDR< 0.01. The overlap of ATAC-seq peak with promoter of genes were defined as peaks within TSS ± 1 kb.

### Transcription factor (TF) binding motif analysis of snATAC-seq data

We determined TF binding motif enrichment in accessible peaks using chromVAR (v.1.4.1)^87^ on either all excitatory neuron population or on cells of individual neuron and non-neuronal cell type/subtypes. GC bias was corrected based on BSgenome.MfascicμLaris.NCBI.5.0 genome by addGCBias function. Then we fetched the human motifs in JASPAR database by getJasparMotifs function. The deviation z-scores for each TF motif in each cell was calculated by computeDeviations function, and high variance TF motifs were obtained by computeVariability function with cut-off at 1.5. For excitatory neurons, Wilcoxon test were performed on TF motif deviation z-scores between 9 subtypes and motifs with pval≤0.01 were determined as differentially enriched motifs. For cells in individual neuron and non-neuronal cell type/subtypes, Wilcoxon test were performed on TF motif deviation z-scores between PFC, M1 and V1 region and motifs with pval ≤ 0.01 were determined as differentially enriched motifs.

To identify the TF binding motif enriched along OLI maturation, we proceeded all peaks of OPC and OLI to compute variability and TF z-scores using chromVAR as described above. Then, we ordered cells by pseudotime described below and kept motifs enriched in all cells.

We used findMotifsGenome.pl tool in HOMER software^88^ to perform TF enrichment analysis on differential accessible peaks of 23 snATAC-seq cell clusters. Then we normalized the results of known motif by p-value and visualized with selected top 30 motifs for each cell cluster. Finally, we used min-max normalization for each TF motif across 23 cell clusters.

### Stereo-seq library preparation and sequencing

Tissues sampled from two female 60-month-old cynomolgus monkeys were embedded with Tissue-Tek OCT (Sakura, 4583) and snap-frozen in liquid nitrogen prechilled isopentane, then transferred to a −80 °C freezer for long time storage. 10 μm cryosections were cut in Leika CM1950 cryostat and placed either in glass slide for Nissl staining or pre-chilled Stereo-seq chips for Stereo-seq procedures. Stereo-seq libraries were prepared as previous describe^49^. Briefly, sections adhered to the Stereo-seq chips were incubated at 37 °C for 3 minutes, then fixed in methanol at −20 °C for 40 minutes. After fixation, the chips with tissues were taken out and dry in the air, permeabilized at 37 °C for 6 min, and reversed transcribed for 2 h at 42 °C. After that, tissue was removed from the chips, then first strand cDNA was released from chips using proteinase K overnight at 55 °C. After purified, the first strands were amplified with KAPA HiFi Hotstart Ready for 15 cycles. The result cDNA was purified and fragmented using in-house Tn5 transposase at 55 °C for 10 min. Fragmentation products were amplified for 13 cycles, then the library was sequenced on DIPSEQ T1 sequencer at China National Gene Bank (CNGB).

### Stereo-seq data processing

The raw data of Stereo-seq were processed as the previous study^49^. Briefly, the identity of coordinates was mapped the designed coordinates matrix of the in situ captured chip which allowing 1 base mismatch and UMIs with N base or more than 2 bases with quality lower than 10 were filtered out. The retained reads were then aligned to the macaca fascicularis genome using STAR, Mapped reads with MAPQ >= 10 were counted and annotated to their corresponding genes using the handleBam^49^ and used to generate a gene expression profile matrix. In this study, we divided the gene expression profile matrix into non-overlapping bins covering an area of 50 × 50 DNB, and the transcripts of the same gene aggregated within each bin. After this step, data were normalized using SCTransform function and unsupervised clustering by Seurat.

To deconvolution of cell types within each bin of Stereo-seq, we first down sampling the cell number of each cell type to 100 in the corresponding snRNA-seq data, and then applied SPOTlight to deconvolute and mapped 20 cell types to bins of Stereo-seq sections. We set probability lower than 0.2 of each cell type as noise, then chose the highest probability out of all cell type as the identified cell type for each bin.

### Inter-regional comparative analysis across cell type/subtypes

We subset the cell type/subtypes of DNBelab C4 RNA-seq data, and compared the differential expressed genes (DEGs) of three regions for each cell type/subtype with R package Seurat (v3.1.1). Cell type/subtypes were subset, followed by Log normalization, and we used the FindAllMarkers function to find the differentially expressed genes in 3 different regions of each cell type/subtype. DEGs of three regions in different cell type/subtype were defined as those with a Foldchange > 1.5, positive ratio over 20% and adjusted P < 0.01. We use the subsetArchRProject function of the R package ArchR (v1.01) to subset the cell type/subtype of snATAC-seq data and compare the differences between the regions within each subtype separately and select DA peaks with a p value < 0.01 and Foldchange > 1.

The Stereo-seq dataset was converted to a Seurat object, differentially expressed genes analysis of three cortical regions for each cell type/subtype were performed with R package Seurat (v3.1.1), followed by similar processing with RNA data. Differentially expressed genes of three regions in each cell type/subtype were defined as those with adjusted P < 0.01.

### Gradient gene expression pattern analysis by Mfuzz

We subset the subtypes of excitatory neuron of DNBelab C4 RNA-seq data, and then calculate the average expression value of all genes in each EX-subtype. Subsequently, the Mfuzz package^47^ was used to characterize the genes with the same gradual expression pattern. For each gene in the average expression matrix, we added 0.000001 to avoid missing value. Then, we filtered genes and standardized the expression using ‘filter.std(min.std=0)’ and ‘standardize()’, and performed M-estimation using “mestimate” function. For the gradient expression pattern across L2/3 IT, L4/5 IT, L5 IT and L6 IT types in each cortical area, the number of clusters was set to 6. For the gradient expression pattern across PFC, M1 and V1 in each EX-subtype, the number of clusters was set to 5. Genes with probability of matching the pattern > 0.5 and maximum expression > 1 were retained for further analysis. Similar clustering analysis by Mfuzz is also applied to Stereo-seq data for the gradient expression pattern across L2/3 IT, L4/5 IT, L5 IT and L6 IT types in each cortical area, genes with probability of matching the pattern > 0.5 were retained for further analysis.

### Pseudotime trajectory analysis of OLI lineage datasets

To uncover the dynamic change of both transcriptome and chromatin accessibility along the oligodendrocyte lineage. We constructed the development trajectory using oligodendrocytes and oligodendrocyte precursor cells. For RNA-seq data, monocle 2 ^89^ was used to order 11706 Oligo cells and 5552 OPCs from the single nuclei RNA-seq data set to developmental trajectory. TPM matrix of OLI cells and OPCs was input and then transformed into normalized mRNA counts using the “relative2abs” function. Genes used for ordering the cells along the trajectory were selected under the following criteria: mean_expression >= 0.5; dispersion_empirical >= 1. After that, the discriminative dimensionality reduction with trees (DDRTree) method was used to reduce data into two dimensions. The first 2000 genes significantly varied across pseudotime were visualized using heatmap and gene ontologies were identified using metascape (https://metascape.org).

For ATAC-seq data, we used monocle 2 to plot a heatmap of gene activity over pseudo time in OPCs differentiation into OLI. We converted the peak by cells matrix to the gene activity score matrix by custom distance-weighted accessibility models in ArchR. To determine peaks that were differentially accessed between the different cell states, we applied differentialGeneTest function from monocle 2 to gene activity score matrix, then used top 1000 genes with the smallest p value to construct a trajectory. DDRTree is used to reduce dimensions and to reconstruct the temporal progression. Finally, we used genes with q-values less than 0.01 to plot heatmap of dynamics genes activity along trajectory. To establish a heatmap of TF motifs enrichment over pseudotime, the cells were grouped into 10 bins based on their trajectory values.

### Evaluation of enrichment of disease risk variants within cell-types

we used the LDSC [https://github.com/bulik/ldsc] to predict enrichments of diseases SNP-heritability in differentially accessible peaks for each cell type or subtype. First, we used the liftOver [https://genome.ucsc.edu/cgi-bin/hgLiftOver] to lift the differentially accessible peaks of each cell type/subtype and all accessible peaks to human hg19 genome. Then, LD scores of SNPs of 1000 genomes phase 3 in those peaks were estimated according to recommended workflow. We generated the summary statistics from previous GWAS studies reported in publications (Supplementary Table 12) and UK Biobank (https://www.ukbiobank.ac.uk/) and calculate the significant enrichment of each diseases following the “cell type specific analysis” workflow recommend by the LDSC authors.

To capture the perturbation of neurology disease in different cortical region of Stereo-seq section, we used a similar analysis as previous study^12^. Briefly, we adapted the homologous in *Macaca fascicularis* genome for up-regulated genes and down-regulated genes patients of ASD, bipolar disorder (BD) and SCZ from recent large scale postmortem brain datasets^69^ (from DGE - Differential Gene Expression), and filtered out the genes with the FDR >0.05 or not captured by the Stereo-seq. Differentially expressed genes of cell type/subtype in each cortical region were calculated by Seurat and using the cutoff of p values < 0.05, then the significance of each disease associated gene set in each cell type/subtype were calculated by Fisher’s exact test, and the *P* values < 0.05 were considered as significant enrichment in the corresponding cell type/subtype.

